# Conformational selection and redox-dependent destabilization in the IspG–FldA electron transfer complex

**DOI:** 10.64898/2026.09.17.752318

**Authors:** Stefan Loonen, Sem Widjaja, Gregory Bokinsky, Nikolina Šoštarić

## Abstract

Electron transfer between proteins is mediated by transient complexes that sample many binding orientations, only a subset productive. Protein electron carriers (PECs) form such complexes with diverse enzymes across central metabolism, yet no general framework predicts which PEC–enzyme pairs will function. A representative case is IspG, the iron–sulfur enzyme catalyzing the penultimate methylerythritol phosphate pathway reaction, whose [4Fe—4S] cluster must be reduced after each turnover by the *Escherichia coli* flavodoxin FldA. It is not known whether FldA engages the open (substrate-free) or closed (substrate-bound) conformation of IspG, or how the cofactor redox states shape their interaction. Using atomistic molecular dynamics with custom cofactor parameters, we find that FldA preferentially engages the open conformation, implying that reduction precedes substrate binding. A positively charged arginine patch anchors FldA, and a single residue, Tyr58, provides a short tunneling bridge to the cluster. Electron transfer then weakens this interface, driving the complex toward dissociation — pointing to an intrinsic mechanism by which a PEC binds its targets transiently yet productively: conformational selection recruits the partner, and electron transfer releases it.

## 1 Introduction

Protein–protein interactions range from stable, high-affinity complexes to transient, low-affinity encounters^1,2^. Stable complexes such as ATP synthase^3^ or the ribosome^4^ adopt well-defined structures, which crystallography and cryo-EM can resolve reliably and that machine-learning methods now increasingly predict from sequence alone^5,6^. Many other functionally important interactions are instead transient by design, allowing a single protein to engage many partners in turn, for example, when a kinase phosphorylates its substrates, a chaperone binds its clients, or a protease cleaves its targets^7,8,9^. Transient complexes are considerably harder to characterize than stable ones^10^, as their weak affinity and short lifetimes keep them from settling into a single, crystallizable conformation. As a result, the machine-learning predictors that now excel on stable complexes are less reliable for transient ones^10,11^. Biological electron transfer often relies on such transient encounters, in which a brief protein contact brings two redox centers (protein-bound cofactors or redox-active residues) close enough for an electron to tunnel from donor to acceptor^12,13^. Rather than locking into one dominant binding mode, the interface samples a range of relative orientations, only a minor subset of which supports productive electron transfer^14^. Conformational transitions into the productive subset act as a gate on the overall reaction^15,16^. However, which orientations support transfer, and how a complex reaches them, remains poorly understood for many electron-transfer pairs.

Protein electron carriers (PECs) such as flavodoxins and ferredoxins serve as electron donors, delivering electrons to a wide range of redox-active enzymes across central metabolism^17,18^ . In metabolic engineering, however, redox-active enzymes expressed in non-native hosts often fail because the host’s available PECs cannot productively reduce the introduced enzyme^20,21^ . The molecular basis of PEC–enzyme compatibility remains unresolved, which limits both the predictive selection and the rational engineering of productive pairs.

A representative example for studying the compatibility of PEC–enzyme pairs is IspG, an iron–sulfur enzyme that catalyzes the penultimate reaction of the methylerythritol phosphate (MEP) pathway, converting the cyclic diphosphate MEcPP into HMBPP^23,24^. The MEP pathway converts central metabolic intermediates into the universal C_5_ precursors of terpenoids^25^, a chemically diverse class of molecules used as materials, fragrances, therapeutic agents, and fuels^26,27^. Catalysis requires the [4Fe–4S] cluster of IspG to be reduced again after each turnover, making the enzyme directly dependent on a compatible electron donor^28,29^. In *E. coli*, this reduction is performed by flavodoxin 1 (FldA), whose low-potential FMN cofactor is one of the few electron carriers thermodynamically competent to reduce the [4Fe–4S] cluster^30,31^. During catalytic turnover, the [4Fe–4S] cluster of IspG cycles between its oxidized (+2) and one-electron-reduced (+1) forms^28^, and the FMN cofactor of FldA cycles between its semiquinone and hydroquinone states^30^. IspG also adopts open (substrate-free) and closed (substrate-bound) conformations^32,33^ .

It remains unknown whether FldA engages the open or the closed conformation of IspG, and how the redox states of the [4Fe–4S] cluster and FMN cofactor shape the FldA–IspG interaction^29^. These conformational and redox states are difficult to isolate experimentally, so we turned to atomistic molecular dynamics, simulating the IspG–FldA complex in three relevant states. IspG’s [4Fe–4S] cluster is ligated by three cysteines, rather than the canonical four, which required deriving quantum-mechanical parameters for the [4Fe–4S] cluster in both oxidation states. The simulations reveal two interconnected layers of selectivity. First, FldA preferentially engages the open, substrate-free conformation of IspG: in the open form an arginine patch on IspG is exposed and pairs electrostatically with three consecutive aspartates (D135–D137) on FldA, whereas substrate binding closes the enzyme and buries the patch. Second, electron transfer itself reshapes the complex — reducing the cluster and oxidizing FMN weakens the protein–protein interface and begins to drive the two proteins apart.

Our results are consistent with the conformational-gating and dynamic-docking frameworks developed for other electron-transfer pairs^15,16^ (see Note S1). They extend these frameworks by showing that the cofactor redox state acts on complex stability: electron transfer weakens the interface that holds the two proteins together. For the IspG–FldA system, this suggests that electron delivery and complex disassembly are linked — the same event that reduces the cluster begins to destabilize the complex, which we propose primes FldA for release and couples electron transfer to the progression of the catalytic cycle. The arginine patch, more tentatively, may help predict whether a given IspG ortholog is compatible with *E. coli* ‘s FldA; whether analogous electrostatic patches are a general feature of PEC–enzyme interfaces, or specific to this pair, remains to be established.

## 2 Results

### 2.1 Both open and closed IspG conformations form complexes with FldA

To test whether FldA engages the open or the closed conformation of IspG, we first built IspG–FldA complex models with IspG in each conformational state.

No experimental structure of *E. coli* IspG is available, so we benchmarked five modeling approaches against the *T. thermophilus* crystal structures of IspG in its open (PDB ID: 2Y0F^33^) and closed (PDB ID: 4G9P^34^) states, which share 38% sequence identity with *E. coli* IspG, with the cluster-coordinating cysteines conserved between the two (Fig. S2).

The open and closed conformations differ by a hinge-like domain closure that brings the [4Fe–4S] cluster proximal to the substrate (Fig. 1C). Four of the five approaches were AI-based structure prediction methods (AlphaFold3, Boltz-2, Chai-1, and Robetta). Because these methods generate a single lowest-energy prediction rather than allowing conformation-specific guidance, they cannot reliably be directed toward a particular conformational state of IspG. The fifth, MODELLER-based homology modeling, reproduced both the open and closed conformations most accurately, as measured by C*α* root mean squared deviation (RMSD) against the *T. thermophilus* reference structures (Fig. 1A), capturing the hinge-like closure at the monomer level. Homodimeric models were assembled by aligning the monomers to the *T. thermophilus* structures (Fig. 1D).

**Figure 1.**
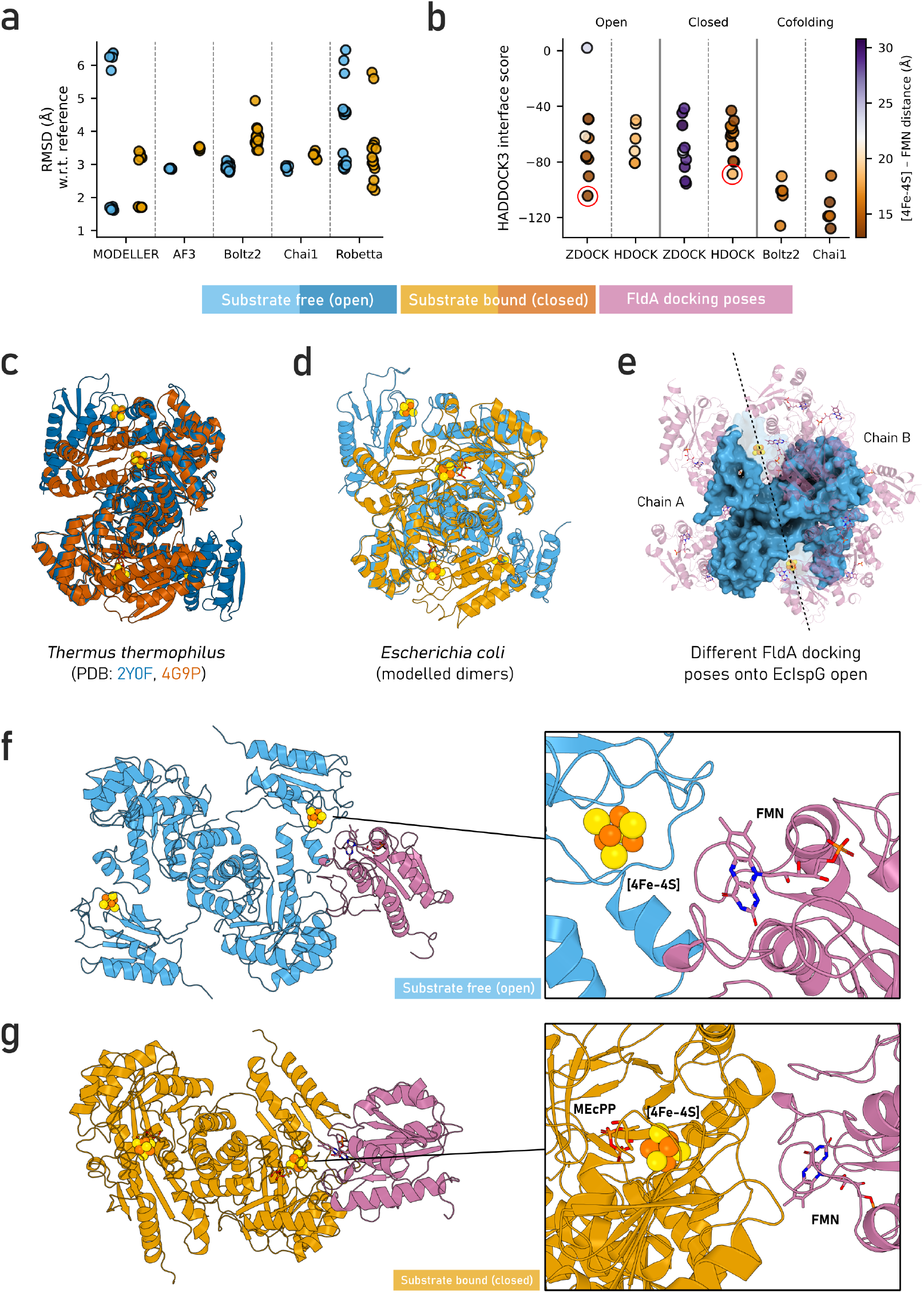
Construction of IspG–FldA complex models in the open and closed conformations of IspG. (A) Comparison of monomer modeling techniques for *E. coli* IspG, evaluated by C*α* RMSD against the *T. thermophilus* crystal structures 2Y0F (substrate-free, blue) and 4G9P (substrate-bound, orange). Homology modeling, implemented through MODELLER, reproduces both conformations most accurately and was selected for downstream steps. (B) HADDOCK3 interface scores for all IspG–FldA docking poses, grouped by method and IspG conformational state (open, closed, or cofolding). Each point represents one pose, colored by the FMN–[4Fe–4S] inter-cofactor distance (Å). Cofolding methods (Boltz-2, Chai-1) were not templated to a specific IspG conformation and are shown as a separate group. The three-step filter (conformation match, inter-cofactor distance below 20 Å, and best HADDOCK3 score) yielded a single selected pose per conformational state, encircled in red. (C) Reference *T. thermophilus* IspG crystal structures showing the substrate-free (open) and substrate-bound (closed) conformations, related by a hinge-like domain closure that positions the [4Fe–4S] cluster proximal to the substrate. (D) *E. coli* IspG homology models in the open and closed states, assembled as homodimers. (E) Representative subset of docking poses of FldA (blue surface) onto the open *E. coli* IspG dimer (pink cartoon), viewed along the homodimer axis (dashed line) that separates chain A (left) and chain B (right). Each chain consists of an N-terminal TIM-barrel domain and a C-terminal cluster-coordinating domain; the [4Fe–4S] cluster is shown in yellow and orange. Poses were chosen to illustrate the spread of predicted FldA binding sites across the IspG surface.(F, G) Final selected models of the IspG–FldA complex in the open (F) and closed (G) conformations, used as starting points for molecular dynamics simulations.

The crystal structure of FldA (PDB ID: 1AHN^35^) was used directly, with the missing C-terminal residues modelled by MODELLER-based loop refinement. Docking it onto each IspG model with two rigid-body methods (ZDOCK, HDOCK) and two cofolding methods (Boltz-2, Chai-1) produced ensembles in which FldA associated at multiple distinct IspG surface regions (Fig. 1E). To select one representative complex per IspG conformational state, we applied a three-step filter: poses were first required to match the intended IspG conformation (open or closed) and to have an inter-cofactor (FMN–[4Fe–4S]) distance below 20 Å. The poses that passed were ranked by HADDOCK3 score (a linear combination of physics-based energy terms and buried surface area; Fig. 1B), and the best-scoring pose per conformation was selected. This yielded a ZDOCK pose for the open state (Fig. 1F) and an HDOCK pose for the closed state (Fig. 1G), which served as the starting points for all subsequent molecular dynamics simulations.

### 2.2 Custom parameters describe the [4Fe– 4S] cluster and its redox-dependent geometry

To simulate the IspG–FldA complex in its different redox states, we first needed an accurate description of both redox cofactors. FldA shuttles electrons through its FMN cofactor, which cycles between oxidized, semiquinone, and hydroquinone states (Fig. 2A). We used the fully reduced hydroquinone (NHQ) and the one-electron-oxidized neutral semiquinone (NSQ), paired with the [4Fe–4S] cluster in the +2 and +1 states, respectively, to represent the preand post-electron-transfer complex.

**Figure 2.**
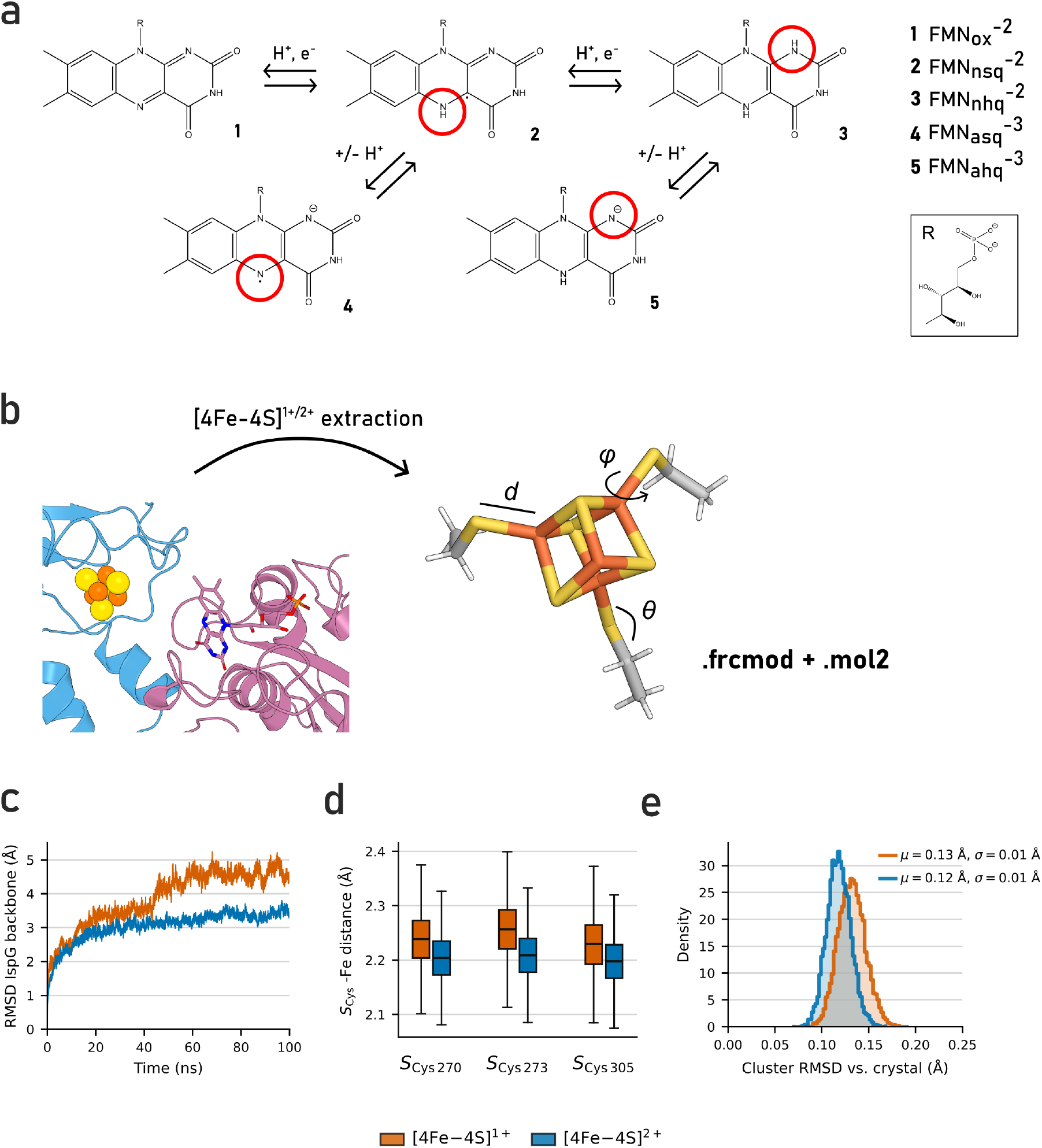
Parameterization and validation of redox-active cofactor force-field parameters. (A) Redox states and interconversions of the FMN cofactor: oxidized (OX), neutral semiquinone (NSQ), neutral hydroquinone (NHQ), anionic semiquinone (ASQ), and anionic hydroquinone (AHQ). Red circles highlight the atom whose protonation or redox state changes at each step. The simulations used the NHQ (two-electron reduced) and NSQ (one-electron reduced) states. Truncated cluster–cysteine QM model used for broken-symmetry DFT parameterization, comprising the [4Fe–4S] core and the side chains of the three coordinating cysteines (Cys270, Cys273, Cys305). Labels indicate the classes of bonded parameter derived from the QM calculation — bond lengths (*d*) and angles (*φ, θ*) — which together with the fitted charges are written to the .frcmod and .mol2 parameter files. (C) Backbone RMSD of the substrate-bound IspG dimer during 100 ns parameter validation simulations in the [4Fe–4S]^+^ and [4Fe–4S]^2+^ states. (D) Distribution of S_Cys_–Fe coordination bond lengths for each coordinating cysteine in both oxidation states; the slight elongation in the reduced state is consistent with experimentally observed redox-induced bond lengthening in [4Fe–4S] clusters. (E) RMSD of the [4Fe–4S] cluster atoms relative to the crystal structure reference (PDB: 4G9P), in both oxidation states.

No existing force field covers IspG’s three-cysteine cluster, in which one iron is left uncoordinated, so we derived a full set of bonded and charge parameters for it from quantum mechanics. A broken-symmetry DFT workflow in MCPB.py (B3LYP, with LANL2DZ on iron and 6-31G* elsewhere) applied to a truncated cluster–cysteine model (Fig. 2B) provided bonded parameters by the Seminario method and RESP (restrained electrostatic potential) charges from the fitted electrostatic potential. For FMN, whose bonded parameters are available through GAFF2, we derived redox-state-specific RESP charges for each relevant state, and applied the same charge-derivation procedure to the substrate of IspG, MEcPP.

To test these parameters, we ran two 100 ns simulations of the substrate-bound IspG dimer, one in each [4Fe– 4S] cluster oxidation state. The IspG backbone RMSD with respect to the initial frame equilibrated within a few nanoseconds, at approximately 3 Å for the +2 state and 4–5 Å for the +1 state, confirming that the overall fold was preserved (Fig. 2C). The Fe–S coordination geometry remained stable, and the parameters reproduced the slight bond elongation expected on one-electron reduction, with S_Cys_–Fe distances shifting from *∼*2.20 Å (2.15–2.25 Å) in the oxidized state to *∼*2.25 Å (2.20–2.30 Å) in the reduced state (Fig. 2D)^36,37^. The cluster core remained close to the crystal reference, with heavy-atom RMSDs of 0.12 *±* 0.01 Å (+2) and 0.13 *±* 0.01 Å (+1) (Fig. 2E).

Together, these results show that the parameters give a stable and physically consistent description of the cluster. Because each oxidation state was parameterized separately, these parameters also reproduce the cluster’s redox-dependent geometry, whereas parameters derived for a single oxidation state and applied to both would fix bond lengths at a single value regardless of redox state. The parameters are made available as Supplementary Data and are transferable to other iron–sulfur enzymes that share this non-canonical three-cysteine coordination environment.

### 2.3 The open conformation of IspG forms an electron transfer-competent complex with FldA

We next asked whether FldA preferentially interacts with the substrate-free (open) or substrate-bound (closed, with MEcPP in the active site) conformation of IspG, simulating both complexes in the pre-electron transfer state, defined by a [4Fe–4S]^2+^ cluster and FMN in the hydroquinone (NHQ) form. To that aim, we compared their electron transfer competence, interface composition, and binding energetics.

Efficient electron transfer between proteins requires the donor and acceptor redox centers to be (i) positioned in close proximity and (ii) well coupled electronically, as measured by the electron tunneling rate. These two requirements are linked through Marcus theory^38,39^: in the nonadiabatic regime that governs inter-protein electron transfer at separations above *∼*10 Å^40,41^, the rate constant is proportional to the square of the electronic coupling, *k*_ET_ *∝* |*H*_DA_|^2^ (full expression in Supplementary Methods). The coupling itself decays exponentially with donor– acceptor distance, but the rate of decay depends on the intervening medium. Within the Pathways formalism^42,43^, per-bond attenuation factors of *ε*_cov_ = 0.6 (covalent bond), *ε*_H_ = 0.36 (hydrogen bond), and *ε*_space_ = 0.5 *e*^*™*1.7(*r™*1.4)^ (through-space jump over a distance *r* in Å) mean that a pathway routed through covalent bonds can be orders of magnitude more conductive than one crossing a poorly packed interface or a vacuum gap.

To assess electron transfer competence of the IspG–FldA complexes, we identified representative donor–acceptor geometries by conformational clustering of each simulation and computed *H*_DA_ between the FMN cofactor of FldA and the [4Fe–4S] cluster of IspG using the Pathways formalism^44^. Because the electron transfer rate is dominated by the highest-coupling conformations sampled across the trajectories rather than by the median, we focus on peak coupling values. These high-coupling conformations are sampled across multiple replicate MD runs and are not confined to sparsely populated conformational clusters (Fig. S4).

The open conformation reaches peak couplings of *∼*10^*™*5^, whereas the closed conformation does not exceed *∼*10^*™*7^ (Fig. 3A). Because *k*_ET_ *∝* |*H*_DA_|^2^, a two-order-of-magnitude difference in coupling corresponds to approximately four orders of magnitude in the maximum accessible electron transfer rate. Without explicit values for Δ*G* and *λ* we cannot determine absolute rates, but the difference between the two conformations is unambiguous: reduction of IspG by FldA proceeds far more efficiently through the open state. Visualizing the dominant tunneling pathways reveals the structural basis of this difference: in the open conformation, the [4Fe–4S] cluster sits in an exposed pocket, so when FldA docks, the donor and acceptor are close enough for short, direct routes. The dominant pathway exits FMN at N5, crosses four backbone atoms of Tyr58 (N, C*α*, C, O), and reaches the non-coordinated Fe4 without leaving FldA (Fig. 3B); at a coupling of *∼*5 *×* 10^*™*5^, it is roughly an order of magnitude above the next-best paths. Still in the open conformation, several secondary pathways in the low 10^*™*6^ range also remain entirely within FldA, routing instead through Tyr96 or via Tyr58 with an additional bridging residue; these contribute meaningfully to the overall coupling distribution despite their lower individual values. In the closed conformation, the cluster is buried beneath IspG, so any pathway must cross the protein–protein interface and traverse multiple IspG residues before reaching it — a structurally longer and electronically less conductive route (Fig. 3C).

**Figure 3.**
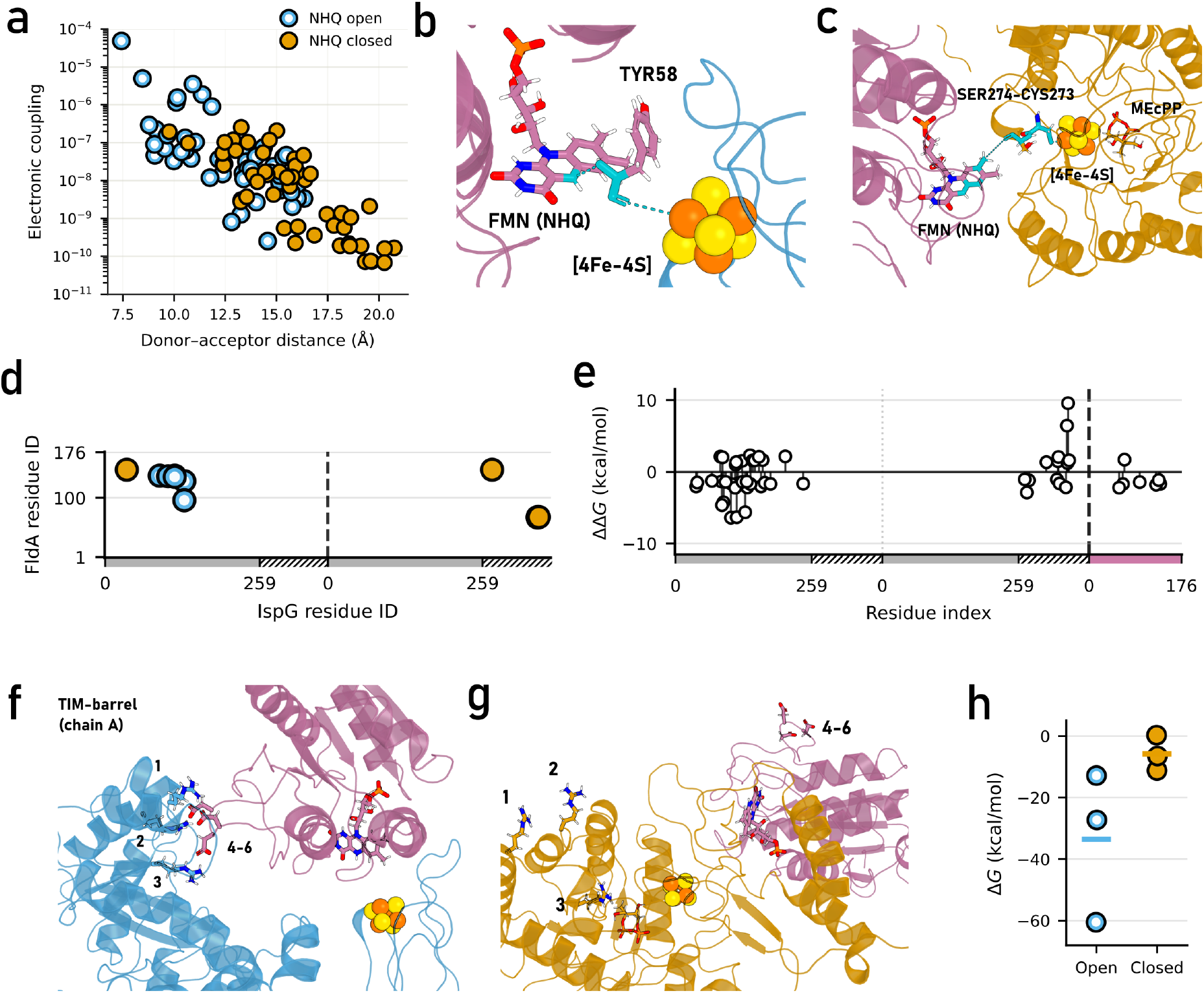
The open conformation of IspG forms an electron transfer-competent complex with FldA. Comparison of substrate-free (open) and substrate-bound (closed) *E. coli* IspG–FldA complexes in the pre-electron-transfer state (NHQ FldA, [4Fe–4S]^2+^). (A) Electronic coupling versus donor–acceptor distance for representative cluster structures from each simulation. The open state (blue/white scatter) samples peak couplings approximately two orders of magnitude above the closed state (orange scatter). (B) Dominant tunneling pathway in the open conformation: FMN N5 *→* Tyr58 backbone *→* IspG cluster Fe4. Residues flanking Tyr58 are hidden for visual clarity; the pathway exits FldA only at the final bond to Fe4. (C) Dominant tunneling pathway in the closed conformation, crossing the FldA–IspG interface via Ser274 and Cys273 before reaching the cluster (see main text for details). (D) Salt-bridge pair map between IspG residues (x-axis) and FldA residues (y-axis), showing contacts with mean occupancy *>*20% averaged across replicates. In the open conformation, Arg92, Arg106, and Arg117 on the TIM-barrel domain of subunit A form salt bridges with Asp135–Asp137 on FldA (residues labeled 1–6 as in panels F and G). In the closed conformation, these contacts are replaced by a salt bridge between Arg351 on the cluster-coordinating domain and Asp67–Asp68 on FldA. The x-axis is colored by domain: TIM-barrel (gray, 1-259) and cluster-coordinating (hatched, 260-372). (E) Per-residue MM/GBSA decomposition of ΔΔ*G* = Δ*G*_open_ *™*Δ*G*_closed_, averaged across replicates, mapped onto IspG residue number. Negative values indicate a stronger contribution to FldA binding in the open conformation; positive values indicate stronger contribution in the closed conformation. The x-axis is colored as in (D), with the addition of pink for FldA. (F, G) Structural renderings of the arginine–aspartate salt bridges in the open (F) and closed (G) conformations. Numbered labels correspond to IspG residues Arg92 (1), Arg117 (2), Arg106 (3) and FldA residues Asp135 (4), Asp136 (5), Asp137 (6). (H) Global MM/GBSA binding free energies for the open and closed complexes; each data point represents the mean Δ*G* of one replicate (*N* = 3).

Beyond the tunneling pathway, the open and closed states also differ markedly in their interfacial contact patterns. We mapped salt bridges with a mean occupancy above 20% (Fig. 3D, Tab. S2). The pair map reveals two distinct interaction patterns. In the open conformation, FldA engages a cluster of arginine residues on the TIM-barrel domain of IspG subunit A — the catalytic domain that houses the substrate-binding site and undergoes closure upon MEcPP binding — most notably Arg92, Arg106, and Arg117, which contact three consecutive aspartates on FldA (Asp135–Asp137). Structural renderings confirm that this positively charged patch is fully accessible in the open state (Fig. 3F): the three arginine side chains project outward and form direct electrostatic contacts across the interface. In the closed conformation, domain closure buries this patch, the contacts with Asp135–Asp137 are lost, and the dominant interactions shift instead to Arg351 on the cluster-coordinating domain of subunit B, which engages Asp67 and Asp68 of FldA (Fig. 3G). Per-residue decomposition of the difference in binding free energy between the two conformations (ΔΔ*G* = Δ*G*_open_ *™*Δ*G*_closed_) confirms that this structural shift has a clear energetic signature: the TIM-barrel domain of subunit A (residue 1-259) is the dominant contributor to binding in the open conformation, whereas the cluster-coordinating domain of subunit B (residue 260-372) takes over in the closed conformation (Fig. 3E). At the global level, MM/GBSA calculations rank FldA binding as more favorable to substrate-free than to substrate-bound IspG, with two of three open replicates below every closed replicate and the third at the closed range’s upper edge (Fig. 3H). The interface also varies less over time in the open conformation, as measured by the RMSD of FldA relative to IspG, indicating that FldA samples a narrower set of bound configurations when IspG is open (Fig. S3B). Furthermore, the IspG backbone itself remained stable across all of these simulations (Fig. S3A), though the TIM-barrel shows slightly lower backbone fluctuations in the closed conformation, consistent with stabilization by substrate binding (Fig. S3C).

Together, these results strongly indicate that FldA preferentially engages the open, substrate-free conformation of IspG. This conformation exposes a positively charged arginine patch on the TIM-barrel domain that anchors FldA across the dimer interface and positions the two cofactors in geometries compatible with efficient electron transfer. Upon substrate binding and domain closure, this patch is obscured, the interface shifts toward the cluster-coordinating domain of the opposing subunit, and the resulting binding mode is both less stable and electronically less competent.

### 2.4 Electron transfer destabilizes the IspG–FldA complex

Having established that FldA preferentially engages the open conformation of IspG, we next asked how the complex responds to electron transfer itself. We compared simulations of the IspG_open_–FldA complex in a pre-transfer state (NHQ FMN, [4Fe–4S]^2+^) and a post-transfer state (NSQ FMN, [4Fe–4S]^1+^).

Across these simulations, global MM/GBSA binding free energies are on average less favorable post-transfer (Fig. 4A), consistent with reduction of the cluster and oxidation of FMN weakening the protein–protein interaction. Per-residue decomposition of ΔΔ*G* = Δ*G*_NHQ_ *™*Δ*G*_NSQ_ maps this change onto the TIM-barrel domain of subunit A, which anchors FldA in the pre-transfer open complex, though the ΔΔ*G* shift is weaker than the domain redistribution seen between the open and closed conformations (Fig. 3E,4B). The change is spread across many residues, and the anchoring interface is broadly preserved: the subunit-A arginine patch (Arg92, Arg106, Arg117, Arg133) that engages Asp135–Asp137, Glu96, and Glu128 of FldA is retained in the post-transfer (NSQ) state, though the relative occupancies of individual salt bridges shift (Tab. S2). The interface therefore weakens overall rather than losing a specific contact.

**Figure 4.**
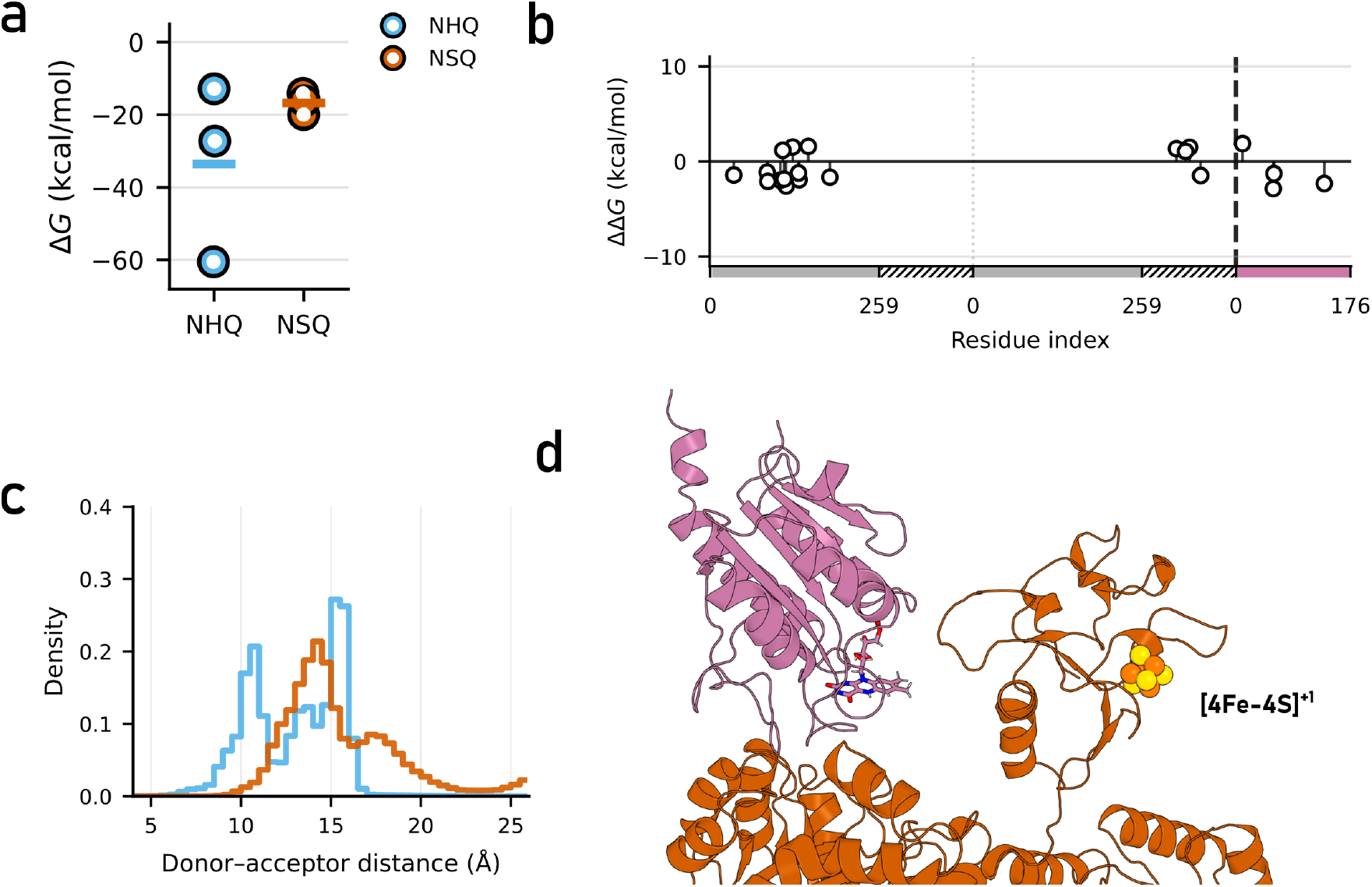
Electron transfer destabilizes the IspG–FldA complex. Comparison of pre-transfer (NHQ FldA, [4Fe–4S]^2+^) and post-transfer (NSQ FldA, [4Fe–4S]^1+^) states of the open *E. coli* IspG–FldA complex. (A) Global MM/GBSA binding free energies for the pre- and post-transfer states; each data point represents the mean Δ*G* of one replicate (*N* = 3). Binding is less favorable on average in the post-transfer state. (B) Per-residue MM/GBSA decomposition of ΔΔ*G* = Δ*G*_NHQ_ *™*Δ*G*_NSQ_, averaged across replicates, mapped onto IspG residue number. Negative values indicate a stronger contribution to FldA binding in the pre-transfer state; positive values indicate stronger contribution in the post-transfer state. The change is localized to the subunit-A TIM-barrel domain but distributed across many residues rather than concentrated in a few. The x-axis is colored by domain: TIM-barrel (gray) and cluster-coordinating (hatched gray). (C) Distribution of the minimum heavy-atom FMN–[4Fe–4S] distance for the pre-transfer (NHQ, blue) and post-transfer (NSQ, orange) states, pooled across replicates. The short (*∼*10 Å) configurations that dominate the pre-transfer ensemble are partly depleted after transfer, shifting the distribution toward larger donor–acceptor separations. (D) Structural rendering of the reoriented [4Fe–4S] cluster in the post-transfer replicate in which the cluster-coordinating domain rotates outward, transiently exposing the reduced cluster to bulk solvent.

This weakening is accompanied by a modest increase in donor–acceptor separation. The distribution of minimum FMN–cluster distances shifts to larger values in the post-transfer state, with the short (*∼*10 Å) configurations being more prevalent in the pre-transfer ensemble than in the post-transfer one (Fig. 4C). The per-replicate time series show that the donor–acceptor distance increases over the trajectory in all three post-transfer replicates – modestly in two and markedly in one – whereas the three pre-transfer replicates hold a stable separation throughout (Fig. S5A,B).

In the replicate with the largest separation increase, the cluster-coordinating domain swings outward and exposes the reduced [4Fe–4S] cluster to bulk solvent (Fig. 4D). An exposed reduced cluster is vulnerable to oxidative attack, which hints at a kinetic tension in the catalytic cycle – the cluster must stay protected long enough for substrate to arrive and catalysis to proceed, before it loses the electron it has just received. As this exposure appeared in only one trajectory, establishing how representative it is will require further simulations.

Throughout these changes the IspG fold itself is preserved: the backbone stays stable across the post-transfer trajectories and the per-residue flexibility barely differs from the pre-transfer state (Fig. S5C,D). Post-transfer trajectories still sample high electronic couplings (*∼*10^*™*5^) that match the pre-transfer state (Fig. S5E) — which does not imply reverse transfer, since electron delivery shifts the cofactor redox potentials against it. The interface therefore loosens without the complex or its transfer-competent arrangement breaking down on the timescales that we simulate here.

Taken together, these results point to a role for electron transfer beyond the transfer event itself. Delivering the electron appears to destabilize the IspG–FldA complex: the donor–acceptor distance drifts upward across all three post-transfer replicates, and binding weakens on average across the same subunit-A interface that anchors the pretransfer state. This pairing suggests a mechanism for how FldA targets and then leaves IspG: tight, geometrically selective binding to the open conformation lets FldA locate IspG and reduce it productively; the destabilization that accompanies electron transfer then releases the electron carrier once its work is done, freeing it for further rounds of reduction. Electron transfer would then serve as an intrinsic release signal that couples the selectivity of the encounter to the timely turnover of the protein electron carrier.

## 3 Discussion

In this work, we combined protein–protein docking, atomistic molecular dynamics simulations with custom quantum-mechanical cofactor parameters, and electron-tunneling pathway analysis to investigate how the conformational state of *E. coli* IspG and the redox state of its cofactors together shape its interaction with FldA. Our results reveal two layers of regulation: (i) a strong conformational preference for the substrate-free, open form of IspG, in which the exposed arginine patch anchors FldA and a short tunneling pathway lying within FldA itself — through Tyr58 — couples the cofactors, and (ii) a redox-dependent weakening of the protein–protein interface following electron delivery, which weakens the complex while leaving its transfer-competent donor–acceptor geometry largely intact.

The point in IspG’s catalytic cycle at which FldA reduces the [4Fe–4S] cluster has never been directly resolved. Earlier proposals have generally assumed that the electron donor engages IspG in its substrate-bound, closed conformation. Rekittke *et al*. argued on structural grounds that a flavodoxin or ferredoxin can be positioned close enough to the buried cluster for effective electron transfer in the closed state^45^, while Wang and Oldfield reached the same ordering on mechanistic grounds, with MEcPP first binding the oxidized ([4Fe–4S]^2+^) cluster and the electron delivered once the cyclodiphosphate ring has opened^23^. Our simulations suggest a reordering of these events, showing that FldA docks preferentially onto the open, substrate-free form of IspG and could therefore reduce the cluster before MEcPP arrives, thereby priming IspG for catalysis. Notably, the same event that reduces the cluster also begins to destabilize the complex, and it does so quickly — the interface loosens within the microsecond trajectories — underscoring how transient the productive encounter is.

Beyond conformational selectivity, the two conformations differ in how directly donor and acceptor can be connected. In the open state, the cluster is exposed enough that a single FldA residue — Tyr58 — bridges FMN to the cluster with no IspG residues in between. The closed state offers no such shortcut: the buried cluster forces the pathway through Ser274 and Cys273. The short Tyr58 bridge is thus specific to the open, substrate-free state and is lost once the enzyme closes upon substrate binding. Electron transfer weakens the open IspG–FldA interface across the same arginine patch that anchored it before transfer, so the event that reduces the cluster also begins to release FldA.

In one post-transfer replicate the cluster-coordinating domain swings outward and transiently exposes the reduced cluster to solvent (for *∼*300 ns) — an observation that bears on two known features of IspG biology. It offers a structural reason for the enzyme’s well-documented oxygen sensitivity^21,46^: a reduced cluster that can leave its shielded pocket is transiently open to oxidative attack, the same vulnerability that hinders IspG expression in aerobic heterologous hosts^21^. It also raises the question of how the cycle commits to catalysis before that electron is lost. At physiological concentrations, MEcPP binding should outpace oxidative damage of the exposed cluster by two to four orders of magnitude (Note S2), so transient exposures like the one seen here need not disrupt catalysis. We therefore hypothesize that substrate arrival is itself the commitment step: MEcPP binding triggers domain closure, which re-buries the cluster and tightens the active site, tying electron delivery to catalysis without the cluster ever needing long-term shielding.

From a biotechnological perspective, these findings highlight two distinct molecular handles for engineering MEP pathway flux in microbial cell factories, including heterologous expression contexts where native electron donor compatibility is a limiting factor. The binding interface — centered on Arg92, Arg106, and Arg117 of the IspG TIM-barrel and Asp135–Asp137 of FldA — provides a target for tuning association affinity, while Tyr58 of FldA, the residue mediating the dominant tunneling pathway, provides a target for tuning electron transfer competence directly. The localization of the critical bridge within FldA rather than IspG is itself practically useful: FldA can be mutated^31^ and tested against wild-type IspG without perturbing the essential MEP pathway *in vivo*. More broadly, other *E. coli* flavodoxins that currently fail to reduce IspG efficiently may lack an equivalent of the Tyr58 bridge, and introducing such a residue at the corresponding position could expand the pool of competent cellular reductants^20^. Experimental validation of the main conclusions remains an important next step. The conformational selectivity hypothesis is most directly testable by measuring FldA binding affinity to substrate-free and substrate-bound IspG separately, for instance using non-cleavable substrate analogs to stably occupy the closed conformation while leaving the protein otherwise unperturbed. The contribution of Tyr58 to electron transfer flux is testable through point mutagenesis, with IspG turnover as a readout, which could be quantified as anaerobic MEcPP-to-HMBPP conversion^28^ using the mutated Tyr58 variant as the sole reductant. Hardest to test is the sequence of events after electron delivery. Our simulations show the interface weakening but not full dissociation, leaving open whether electron transfer alone drives FldA off or whether substrate binding completes the displacement — and, relatedly, whether FldA lingers at all once it has delivered the electron, given that in the post-transfer simulations it no longer shields the cluster. Distinguishing these possibilities is difficult because the post-transfer state is a transient catalytic intermediate rather than a species that can be isolated at equilibrium.

Next to the specific IspG–FldA system, this work offers several methodological contributions of broader utility. The derived force-field parameters for the [4Fe–4S] cluster, covering both the +1 and +2 oxidation states, fill a gap not covered by existing parameterizations, namely a coordination environment in which one iron site lacks a protein ligand. These parameters are directly transferable to simulations of other Fe–S enzymes in analogous environments. The redox-state-specific RESP charges derived for FMN in combination with AMBER force fields provide a consistent description of all catalytically relevant FMN redox states, including the semiquinone radical, and serve as a resource for the broader flavoprotein simulation community.

These results also connect to the broader problem raised at the outset: the molecular basis of compatibility between protein electron carriers and their enzyme partners. FldA itself is a promiscuous electron donor that, in *E. coli*, reduces several enzymes beyond IspG — including methionine synthase and biotin synthase^17^. How such a carrier achieves selectivity for productive partners against a background of off-target encounters, and why some carrier–enzyme pairs fail when transplanted into heterologous hosts, remain open questions. For the FldA–IspG case, our findings suggest that a productive encounter requires the alignment of several conditions: an open IspG conformation, engagement of the arginine patch, and a precise geometry that positions Tyr58 of FldA adjacent to the cluster. Electron transfer then weakens this same interface, so the features that select a productive partner also set up the carrier’s release once transfer is complete. Exploring whether this multi-criteria selection logic generalizes to FldA’s other partners, and to promiscuous electron carriers more broadly, is a natural extension of the framework developed here.

More fundamentally, transient electron-transfer complexes of this kind remain invisible to the structureprediction tools that have transformed stable-complex biology. Their functional interface is defined by a conformation- and redox-dependent ensemble, not a single predictable structure. Physics-based simulations are able to capture that ensemble, and with it the selection logic that a single predicted structure would miss. Applying that approach systematically across carrier–enzyme pairs would clarify how productive interfaces are selected from a background of binding events, and would provide the rational basis for compatibility prediction that metabolic engineering currently lacks.

## 4 Methods

A flowchart of the full computational workflow — from IspG modeling and docking through cofactor parameterization, molecular dynamics, and trajectory analysis — is shown in Fig. S1, with full details given in the Supplementary Methods.

### 4.1 System construction and docking

Monomeric *E. coli* IspG models were generated using AlphaFold3^6^, Boltz-2^47^, Chai-1^48^, Robetta^49,50^, and homology modeling with MODELLER^51^. Boltz-2 and Robetta were additionally run using the *T. thermophilus* IspG crystal structures 2Y0F (substrate-free)^33^ and 4G9P (substrate-bound)^34^ as templates. All *E. coli* IspG models were evaluated by C*α* RMSD to both *T. thermophilus* reference structures using CE-align, as implemented in PyMOL 3^52^. Homology modeling reproduced both conformations most accurately and was selected for all subsequent steps. Substrate-free (open) and substrate-bound (closed) IspG monomers were built against the 2Y0F and 4G9P templates, respectively, and assembled into homodimers by aligning two copies of each monomer to the corresponding crystallographic dimer. The [4Fe–4S] clusters and the substrate MEcPP were placed based on the same crystallographic entries. The FldA crystal structure (PDB: 1AHN) was used, for which the experimentally unresolved C-terminal tail was rebuilt using MODELLER.

Rigid-body docking of FldA onto IspG was performed with ZDOCK^53^ and HDOCK^54^ for both IspG conformations, generating 10 poses for ZDOCK and 100 poses for HDOCK per conformation. Cofolding-based docking was additionally performed with Boltz-2 and Chai-1, yielding 5 poses each. All poses were filtered by (i) IspG conformational state, (ii) inter-cofactor distance below 20 Å, and (iii) HADDOCK3^55^ score; the best-scoring pose to pass filter (ii) was selected per conformational state.

### 4.2 Cofactor parameterization

Force-field parameters for the [4Fe–4S] cluster were derived from broken-symmetry DFT calculations^56^ (UB3LYP^57,58^/6-31G*^59,60^/LANL2DZ^61,62,63^) using the Seminario^64^ method as implemented in MCPB.py^65^, for both the oxidized (+2) and one-electron-reduced (+1) redox states of the [4Fe–4S] cluster. RESP (restrained electrostatic potential) charges^66^ were fitted on the optimized cluster model, after which the obtained iron, bridging sulfide, and Cys C*β*/S*γ* atom charges were combined with the standard ff19SB^67^ CYX backbone charges, yielding a patched cluster–cysteine unit. RESP charges were also derived for all five FMN redox states (Fig. 2A)^68^, and for MEcPP at the HF/6-31G* level, following B3LYP/6-31G* geometry optimization. Bonded parameters were obtained using GAFF2^69^ atom types. The fully reduced neutral hydroquinone (NHQ) and the one-electron-oxidized neutral semiquinone (NSQ) FMN states were selected for classical molecular dynamics simulations based on their catalytic relevance.

### 4.3 Molecular dynamics simulations

Molecular dynamics simulations were performed using GROMACS 2024.3^70^ with the AMBER ff19SB force field, the OPC water model^71^, and the custom parameters described above. Protonation states of titratable residues were assigned at pH 7.0 with PDB2PQR^72^ (Amber-Tools25^73^). Each system was solvated in an OPC water box extending 10 Å beyond the protein and neutralized with NaCl. Three conditions of *E. coli* IspG in complex with FldA were simulated (Table 1), chosen to isolate the conformational and redox variables in turn: NHQ FldA with substrate-free (open) IspG, NHQ FldA with substrate-bound (closed) IspG, and NSQ FldA with substrate-free IspG. In the substrate-bound condition, the MEP intermediate MEcPP was present in the IspG active site. Each condition was simulated in triplicate with the same starting structure but different initial velocities, yielding nine independent 1 *µ*s trajectories (9 *µ*s aggregate). Production runs were carried out in the NPT ensemble at 310 K and 1 bar with a 2 fs timestep. Two additional 100 ns trajectories of the substrate-bound dimer were generated to validate the [4Fe–4S] parameters in both redox states of the cluster.

**Table 1.** Redox, conformational, and protonation states of the *E. coli* IspG–FldA system investigated with classical molecular dynamics simulations.

| FMN state | [4Fe–4S] charge | IspG conformation |
| --- | --- | --- |
| NHQ | +2 | Open |
| NHQ | +2 | Closed |
| NSQ | +1 | Open |

### 4.4 Trajectory analysis

Standard structural metrics (RMSD, RMSF, radius of gyration, FMN–[4Fe–4S] cluster minimum distance) were computed with GROMACS built-in tools. IspG–FldA interface contacts, salt bridges, hydrogen bonds and hydrophobic contacts were quantified with a custom MDAnalysis-based^74^ script. Binding free energies were estimated by MM/GBSA^75,76^ using gmx_MMPBSA v1.6.3^77,78^. Representative structures of each trajectory were obtained through GROMOS conformational clustering^79^. Electron transfer pathways between the FMN N5 atom (donor) and the non-coordinated Fe4 of the [4Fe–4S] cluster (acceptor) were evaluated using the VMD Pathways plugin^44^.

## Code and data availability

Force-field parameters for the [4Fe–4S] cluster (in the

+1 and +2 oxidation states), RESP-derived partial charges for the five FMN redox states (OX, NSQ, ASQ, NHQ, AHQ) and for the MEcPP substrate, all GROMACS input files used for production runs and parameter validation, and the Python analysis scripts used to compute interface contacts, salt-bridge occupancies, conformational clustering, electronic coupling distributions, and tunneling pathway statistics are available at https://github.com/StefanLoonen1/ET-in-weak-PPIs and archived at 4TU under DOI: 10.4121/972d06c3-9cde-4e82-bd8e-e15624fc3e09. The QM input and output files used in the broken-symmetry DFT parameterization of the [4Fe–4S] cluster and the RESP charge derivations for FMN and MEcPP are included in the same repository. Production trajectories (9 *µ*s aggregate) are available from the corresponding author on reasonable request due to their large cumulative size.

## Competing interests

The authors declare no competing interests.

## Acknowledgments

We thank P. Navarčíková and A. Pérez de Alba Ortíz, for carefully reading the report and providing valuable feedback. We thank SURF (www.surf.nl) for the support in using the National Supercomputer Snellius, using grant no. EINF-16894 and EINF-18664, as well as the Delft High Performance Computing Center for the use of the DelftBlue Supercomputer

## Notes

https://github.com/StefanLoonen1/ET-in-weak-PPIs

https://doi.org/10.4121/972d06c3-9cde-4e82-bd8e-e15624fc3e09

### Competing Interest Statement

The authors have declared no competing interest.

## References

[1] M. Nooren and J. M. Thornton, “Diversity of protein–protein interactions,” en, The EMBO Journal, vol. 22, no. 14, pp. 3486–3492, Jul. 2003. doi: 10.1093/emboj/cdg359.

[2] J. Mintseris and Z. Weng, “Structure, function, and evolution of transient and obligate protein–protein interactions,” en, Proceedings of the National Academy of Sciences, vol. 102, no. 31, pp. 10930–10935, Aug. 2005. doi: 10.1073/pnas.0502667102.

[3] Y. Lai et al., “Structure of the human ATP synthase,” en, Molecular Cell, vol. 83, no. 12, 2137–2147.e4, Jun. 2023. doi: 10.1016/j.molcel.2023.04.029.

[4] A. Doudna and V. L. Rath, “Structure and Function of the Eukaryotic Ribosome,” en, Cell, vol. 109, no. 2, pp. 153–156, Apr. 2002. doi: 10.1016/S0092-8674(02)00725-0.

[5] J. Jumper et al., “Highly accurate protein structure prediction with AlphaFold,” en, Nature, vol. 596, no. 7873, pp. 583–589, Aug. 2021, Number: 7873. doi: 10.1038/s41586-021-03819-2.

[6] J. Abramson et al., “Accurate structure prediction of biomolecular interactions with AlphaFold 3,” en, Nature, vol. 630, no. 8016, pp. 493–500, Jun. 2024. doi: 10.1038/s41586-024-07487-w.

[7] F. U. Hartl, A. Bracher, and M. Hayer-Hartl, “Molecular chaperones in protein folding and proteostasis,” en, Nature, vol. 475, no. 7356, pp. 324–332, Jul. 2011. doi: 10.1038/nature10317.

[8] R. T. Sauer and T. A. Baker, “AAA+ Proteases: ATP-Fueled Machines of Protein Destruction,” en, Annual Review of Biochemistry, vol. 80, no. 1, pp. 587–612, Jul. 2011. doi: 10.1146/annurevbiochem-060408-172623.

[9] A. Cumberworth, G. Lamour, M. M. Babu, and J. Gsponer, “Promiscuity as a functional trait: Intrinsically disordered regions as central players of interactomes,” en, Biochemical Journal, vol. 454, no. 3, pp. 361–369, Sep. 2013. doi: 10.1042/BJ20130545.

[10] H. Elhabashy, F. Merino, V. Alva, O. Kohlbacher, and A. N. Lupas, “Exploring protein-protein interactions at the proteome level,” en, Structure, vol. 30, no. 4, pp. 462–475, Apr. 2022. doi: 10.1016/j.str.2022.02.004.

[11] R. Yin, B. Y. Feng, A. Varshney, and B. G. Pierce, “Benchmarking <span style=“font-variant:smallcaps;”>AlphaFold</span> for protein complex modeling reveals accuracy determinants,” en, Protein Science, vol. 31, no. 8, e4379, Aug. 2022. doi: 10.1002/pro.4379.

[12] M. Ubbink, “Complexes of Photosynthetic Redox Proteins Studied by NMR,” en, Photosynthesis Research, vol. 81, no. 3, pp. 277–287, Sep. 2004. doi: 10.1023/B:PRES.0000036880.67124.e7.

[13] M. Ubbink, “The courtship of proteins: Understanding the encounter complex,” en, FEBS Letters, vol. 583, no. 7, pp. 1060–1066, Apr. 2009. doi: 10.1016/j.febslet.2009.02.046.

[14] Q. Bashir, S. Scanu, and M. Ubbink, “Dynamics in electron transfer protein complexes,” en, The FEBS Journal, vol. 278, no. 9, pp. 1391–1400, May 2011. doi: 10.1111/j.1742-4658.2011.08062.x.

[15] V. L. Davidson, “Protein Control of True, Gated, and Coupled Electron Transfer Reactions,” en, Accounts of Chemical Research, vol. 41, no. 6, pp. 730–738, Jun. 2008. doi: 10.1021/ar700252c.

[16] Z.-X. Liang et al., “Dynamic Docking and Electron Transfer between Zn-myoglobin and Cytochrome b5,” en, Journal of the American Chemical Society, vol. 124, no. 24, pp. 6849–6859, Jun. 2002. doi: 10.1021/ja0127032.

[17] J. Sancho, “Flavodoxins: Sequence, folding, binding, function and beyond,” en, Cellular and Molecular Life Sciences, vol. 63, no. 7-8, pp. 855–864, Apr. 2006, Number: 7-8. doi: 10.1007/s00018-005-5514-4.

[18] J. T. Atkinson, I. Campbell, G. N. Bennett, and J. J. Silberg, “Cellular Assays for Ferredoxins: A Strategy for Understanding Electron Flow through Protein Carriers That Link Metabolic Pathways,” en, Biochemistry, vol. 55, no. 51, pp. 7047–7064, Dec. 2016, Number: 51. doi: 10.1021/acs.biochem.6b00831.

[19] J. Campbell, G. N. Bennett, and J. J. Silberg, “Evolutionary Relationships Between Low Potential Ferredoxin and Flavodoxin Electron Carriers,” en, Frontiers in Energy Research, vol. 7, p. 79, Aug. 2019. doi: 10.3389/fenrg.2019.00079.

[20] F. D’Angelo et al., “Cellular assays identify barriers impeding iron-sulfur enzyme activity in a non-native prokaryotic host,” en, eLife, vol. 11, e70936, Mar. 2022. doi: 10.7554/eLife.70936.

[21] H. Shomar and G. Bokinsky, “Harnessing iron-sulfur enzymes for synthetic biology,” en, Biochimica et Biophysica Acta (BBA) -Molecular Cell Research, vol. 1871, no. 5, p. 119718, Jun. 2024, Number: 5. doi: 10.1016/j.bbamcr.2024.119718.

[22] H. Shomar and G. Bokinsky, “Towards a Synthetic Biology Toolset for Metallocluster Enzymes in Biosynthetic Pathways: What We Know and What We Need,” en, Molecules, vol. 26, no. 22, p. 6930, Nov. 2021, Number: 22. doi: 10.3390/molecules26226930.

[23] W. Wang and E. Oldfield, “Bioorganometallic Chemistry with IspG and IspH: Structure, Function, and Inhibition of the [Fe4 S4 ] Proteins Involved in Isoprenoid Biosynthesis,” en, Angewandte Chemie International Edition, vol. 53, no. 17, pp. 4294–4310, Apr. 2014, Number: 17. doi: 10.1002/anie.201306712.

[24] J. Perez-Gil, J. Behrendorff, A. Douw, and C. E. Vickers, “The methylerythritol phosphate pathway as an oxidative stress sense and response system,” en, Nature Communications, vol. 15, no. 1, p. 5303, Jun. 2024, Number: 1. doi: 10.1038/s41467-024-49483-8.

[25] J. Lombard and D. Moreira, “Origins and Early Evolution of the Mevalonate Pathway of Isoprenoid Biosynthesis in the Three Domains of Life,” en, Molecular Biology and Evolution, vol. 28, no. 1, pp. 87–99, Jan. 2011. doi: 10.1093/molbev/msq177.

[26] J. Hanssens, D. Meneses, J. M. Saya, and R. V. A. Orru, “Terpenes and Terpenoids: How can we use them?” en, European Journal of Organic Chemistry, vol. 28, no. 24, e202401151, Jun. 2025. doi: 10.1002/ejoc.202401151.

[27] P. K. Ajikumar, K. Tyo, S. Carlsen, O. Mucha, T. H. Phon, and G. Stephanopoulos, “Terpenoids: Opportunities for Biosynthesis of Natural Product Drugs Using Engineered Microorganisms,” en, Molecular Pharmaceutics, vol. 5, no. 2, pp. 167–190, Apr. 2008. doi: 10.1021/mp700151b.

[28] Y. Xiao, G. Zahariou, Y. Sanakis, and P. Liu, “IspG Enzyme Activity in the Deoxyxylulose Phosphate Pathway: Roles of the Iron-Sulfur Cluster,” en, Biochemistry, vol. 48, no. 44, pp. 10483–10485, Nov. 2009, Number: 44. doi: 10.1021/bi901519q.

[29] A. Douw, J. Perez-Gil, G. Schenk, and C. E. Vickers, “Iron–Sulfur Cluster Enzymes of the Methylerythritol Phosphate Pathway: IspG and IspH,” en, Biochemistry, vol. 64, no. 12, pp. 2544–2555, Jun. 2025. doi: 10.1021/acs.biochem.4c00714.

[30] H. Vetter and J. Knappe, “Flavodoxin and Ferredoxm of Escherichia coli,” en, Hoppe-Seyler′s Zeitschrift für physiologische Chemie, vol. 352, no. 1, pp. 433–446, Jan. 1971. doi: 10.1515/bchm2.1971.352.1.433.

[31] K.-J. Puan, H. Wang, T. Dairi, T. Kuzuyama, and C. T. Morita, “fldA is an essential gene required in the 2-C -methyl-<span style=“font-variant:smallcaps;”>D</span> -erythritol 4-phosphate pathway for isoprenoid biosynthesis,” en, FEBS Letters, vol. 579, no. 17, pp. 3802–3806, Jul. 2005. doi: 10.1016/j.febslet.2005.05.047.

[32] M. Lee et al., “Biosynthesis of Isoprenoids: Crystal Structure of the [4Fe–4S] Cluster Protein IspG,” en, Journal of Molecular Biology, vol. 404, no. 4, pp. 600–610, Dec. 2010. doi: 10.1016/j.jmb.2010.09.050.

[33] I. Rekittke et al., “Structure of the E -1-hydroxy-2-methyl-but-2-enyl-4-diphosphate synthase (GcpE) from Thermus thermophilus,” en, FEBS Letters, vol. 585, no. 3, pp. 447–451, Feb. 2011. doi: 10.1016/j.febslet.2010.12.012.

[34] I. Rekittke, H. Jomaa, and U. Ermler, “Structure of the GcpE (IspG)–MEcPP complex from Thermus thermophilus,” en, FEBS Letters, vol. 586, no. 19, pp. 3452–3457, Sep. 2012. doi: 10.1016/j.febslet.2012.07.070.

[35] D. M. Hoover and M. L. Ludwig, “A flavodoxin that is required for enzyme activation: The structure of oxidized flavodoxin from Escherichia coli at 1.8 å resolution,” en, Protein Science, vol. 6, no. 12, pp. 2525–2537, Dec. 1997. doi: 10.1002/pro.5560061205.

[36] H. Beinert, R. H. Holm, and E. Münck, “Iron-Sulfur Clusters: Nature’s Modular, Multipurpose Structures,” en, Science, vol. 277, no. 5326, pp. 653–659, Aug. 1997, Number: 5326. doi: 10.1126/science.277.5326.653.

[37] R. A. Torres, T. Lovell, L. Noodleman, and D. A. Case, “Density Functional and Reduction Potential Calculations of Fe4 S4 Clusters,” en, Journal of the American Chemical Society, vol. 125, no. 7, pp. 1923–1936, Feb. 2003. doi: 10.1021/ja0211104.

[38] R. Marcus and N. Sutin, “Electron transfers in chemistry and biology,” en, Biochimica et Biophysica Acta (BBA) -Reviews on Bioenergetics, vol. 811, no. 3, pp. 265–322, Aug. 1985. doi: 10.1016/0304-4173(85)90014-X.

[39] R. A. Marcus, “Electron transfer reactions in chemistry. Theory and experiment,” en, Reviews of Modern Physics, vol. 65, no. 3, pp. 599–610, Jul. 1993. doi: 10.1103/RevModPhys.65.599.

[40] J. Hopfield, “Electron Transfer Between Biological Molecules by Thermally Activated Tunneling,” en, Proceedings of the National Academy of Sciences, vol. 71, no. 9, pp. 3640–3644, Sep. 1974. doi: 10.1073/pnas.71.9.3640.

[41] R. Winkler and H. B. Gray, “Electron Flow through Metalloproteins,” en, Chemical Reviews, vol. 114, no. 7, pp. 3369–3380, Apr. 2014, Number: 7. doi: 10.1021/cr4004715.

[42] D. N. Beratan, J. N. Onuchic, J. R. Winkler, and H. B. Gray, “Electron-Tunneling Pathways in Proteins,” en, Science, vol. 258, no. 5089, pp. 1740–1741, Dec. 1992. doi: 10.1126/science.1334572.

[43] N. Onuchic, D. N. Beratan, J. R. Winkler, and H. B. Gray, “Pathway Analysis of Protein Electron-Transfer Reactions,” en, Annual Review of Biophysics and Biomolecular Structure, vol. 21, no. 1, pp. 349–377, Jun. 1992. doi: 10.1146/annurev.bb.21.060192.002025.

[44] A. Balabin, X. Hu, and D. N. Beratan, “Exploring biological electron transfer pathway dynamics with the Pathways Plugin for VMD,” en, Journal of Computational Chemistry, vol. 33, no. 8, pp. 906–910, Mar. 2012. doi: 10.1002/jcc.22927.

[45] I. Rekittke, H. Jomaa, and U. Ermler, “Structure of the GcpE (IspG)–MEcPP complex from Thermus thermophilus,” en, FEBS Letters, vol. 586, no. 19, pp. 3452–3457, Sep. 2012. doi: 10.1016/j.febslet.2012.07.070.

[46] M. Seemann et al., “Isoprenoid biosynthesis in chloroplasts via the methylerythritol phosphate pathway: The (E)-4-hydroxy-3-methylbut-2-enyl diphosphate synthase (GcpE) from Arabidopsis thaliana is a [4Fe?4S] protein,” en, JBIC Journal of Biological Inorganic Chemistry, vol. 10, no. 2, pp. 131–137, Mar. 2005. doi: 10.1007/s00775-004-0619-z.

[47] S. Passaro et al., Boltz-2: Towards Accurate and Efficient Binding Affinity Prediction, en, Jun. 2025. doi: 10.1101/2025.06.14.659707.

[48] Chai Discovery et al., Chai-1: Decoding the molecular interactions of life, en, Oct. 2024. doi: 10.1101/2024.10.10.615955.

[49] Y. Song et al., “High-Resolution Comparative Modeling with RosettaCM,” en, Structure, vol. 21, no. 10, pp. 1735–1742, Oct. 2013. doi: 10.1016/j.str.2013.08.005.

[50] M. Baek et al., “Accurate prediction of protein structures and interactions using a three-track neural network,” en, Science, vol. 373, no. 6557, pp. 871–876, Aug. 2021. doi: 10.1126/science.abj8754.

[51] B. Webb and A. Sali, “Comparative Protein Structure Modeling Using MODELLER,” en, Current Protocols in Bioinformatics, vol. 54, no. 1, Jun. 2016. doi: 10.1002/cpbi.3.

[52] Schrödinger, LLC, “The PyMOL Molecular Graphics System, Version 3.0,” May 2024.

[53] B. G. Pierce, Y. Hourai, and Z. Weng, “Accelerating Protein Docking in ZDOCK Using an Advanced 3D Convolution Library,” en, PLoS ONE, vol. 6, no. 9, O. Keskin, Ed., e24657, Sep. 2011. doi: 10.1371/journal.pone.0024657.

[54] Y. Yan, H. Tao, J. He, and S.-Y. Huang, “The HDOCK server for integrated protein–protein docking,” en, Nature Protocols, vol. 15, no. 5, pp. 1829–1852, May 2020. doi: 10.1038/s41596-020-0312-x.

[55] M. Giulini et al., “HADDOCK3: A Modular and Versatile Platform for Integrative Modeling of Biomolecular Complexes,” en, Journal of Chemical Information and Modeling, vol. 65, no. 13, pp. 7315–7324, Jul. 2025. doi: 10.1021/acs.jcim.5c00969.

[56] L. Noodleman, “Valence bond description of antiferromagnetic coupling in transition metal dimers,” en, The Journal of Chemical Physics, vol. 74, no. 10, pp. 5737–5743, May 1981. doi: 10.1063/1.440939.

[57] A. D. Becke, “Density-functional thermochemistry. III. The role of exact exchange,” en, The Journal of Chemical Physics, vol. 98, no. 7, pp. 5648–5652, Apr. 1993. doi: 10.1063/1.464913.

[58] C. Lee, W. Yang, and R. G. Parr, “Development of the Colle-Salvetti correlation-energy formula into a functional of the electron density,” en, Physical Review B, vol. 37, no. 2, pp. 785–789, Jan. 1988. doi: 10.1103/PhysRevB.37.785.

[59] W. J. Hehre, R. Ditchfield, and J. A. Pople, “Self—Consistent Molecular Orbital Methods. XII. Further Extensions of Gaussian—Type Basis Sets for Use in Molecular Orbital Studies of Organic Molecules,” en, The Journal of Chemical Physics, vol. 56, no. 5, pp. 2257–2261, Mar. 1972. doi: 10.1063/1.1677527.

[60] C. Hariharan and J. A. Pople, “The influence of polarization functions on molecular orbital hydrogenation energies,” en, Theoretica Chimica Acta, vol. 28, no. 3, pp. 213–222, 1973. doi: 10.1007/BF00533485.

[61] J. Hay and W. R. Wadt, “Ab initio effective core potentials for molecular calculations. Potentials for the transition metal atoms Sc to Hg,” en, The Journal of Chemical Physics, vol. 82, no. 1, pp. 270–283, Jan. 1985. doi: 10.1063/1.448799.

[62] J. Hay and W. R. Wadt, “Ab initio effective core potentials for molecular calculations. Potentials for K to Au including the outermost core orbitals,” en, The Journal of Chemical Physics, vol. 82, no. 1, pp. 299–310, Jan. 1985. doi: 10.1063/1.448975.

[63] W. R. Wadt and P. J. Hay, “Ab initio effective core potentials for molecular calculations. Potentials for main group elements Na to Bi,” en, The Journal of Chemical Physics, vol. 82, no. 1, pp. 284–298, Jan. 1985. doi: 10.1063/1.448800.

[64] M. Seminario, “Calculation of intramolecular force fields from second-derivative tensors,” en, International Journal of Quantum Chemistry, vol. 60, no. 7, pp. 1271–1277, 1996. doi: 10.1002/(SICI)1097-461X(1996)60:7<1271::AID-QUA8>3.0.CO;2-W.

[65] P. Li and K. M. Merz, “MCPB.py: A Python Based Metal Center Parameter Builder,” en, Journal of Chemical Information and Modeling, vol. 56, no. 4, pp. 599–604, Apr. 2016, Number: 4. doi: 10.1021/acs.jcim.5b00674.

[66] C. I. Bayly, P. Cieplak, W. Cornell, and P. A. Kollman, “A well-behaved electrostatic potential based method using charge restraints for deriving atomic charges: The RESP model,” en, The Journal of Physical Chemistry, vol. 97, no. 40, pp. 10269–10280, Oct. 1993. doi: 10.1021/j100142a004.

[67] C. Tian et al., “ff19SB: Amino-Acid-Specific Protein Backbone Parameters Trained against Quantum Mechanics Energy Surfaces in Solution,” en, Journal of Chemical Theory and Computation, vol. 16, no. 1, pp. 528–552, Jan. 2020. doi: 10.1021/acs.jctc.9b00591.

[68] K. Kar, A.-F. Miller, and M.-A. Mroginski, “Understanding flavin electronic structure and spectra,” en, WIREs Computational Molecular Science, vol. 12, no. 2, e1541, Mar. 2022. doi: 10.1002/wcms.1541.

[69] X. He, V. H. Man, W. Yang, T.-S. Lee, and J. Wang, “A fast and high-quality charge model for the next generation general AMBER force field,” en, The Journal of Chemical Physics, vol. 153, no. 11, p. 114502, Sep. 2020. doi: 10.1063/5.0019056.

[70] J. Abraham et al., “GROMACS: High performance molecular simulations through multi-level parallelism from laptops to supercomputers,” en, SoftwareX, vol. 1-2, pp. 19–25, Sep. 2015. doi: 10.1016/j.softx.2015.06.001.

[71] S. Izadi, R. Anandakrishnan, and A. V. Onufriev, “Building Water Models: A Different Approach,” en, The Journal of Physical Chemistry Letters, vol. 5, no. 21, pp. 3863–3871, Nov. 2014. doi: 10.1021/jz501780a.

[72] E. Jurrus et al., “Improvements to the <span style=“font-variant:small-caps;”>APBS</span> biomolecular solvation software suite,” en, Protein Science, vol. 27, no. 1, pp. 112–128, Jan. 2018. doi: 10.1002/pro.3280.

[73] D. A. Case et al., “AmberTools,” en, Journal of Chemical Information and Modeling, vol. 63, no. 20, pp. 6183–6191, Oct. 2023. doi: 10.1021/acs.jcim.3c01153.

[74] N. Michaud-Agrawal, E. J. Denning, T. B. Woolf, and O. Beckstein, “MDAnalysis: A toolkit for the analysis of molecular dynamics simulations,” en, Journal of Computational Chemistry, vol. 32, no. 10, pp. 2319–2327, Jul. 2011. doi: 10.1002/jcc.21787.

[75] P. A. Kollman et al., “Calculating Structures and Free Energies of Complex Molecules: Combining Molecular Mechanics and Continuum Models,” en, Accounts of Chemical Research, vol. 33, no. 12, pp. 889–897, Dec. 2000. doi: 10.1021/ar000033j.

[76] S. Genheden and U. Ryde, “The MM/PBSA and MM/GBSA methods to estimate ligand-binding affinities,” en, Expert Opinion on Drug Discovery, vol. 10, no. 5, pp. 449–461, May 2015, Number: 5. doi: 10.1517/17460441.2015.1032936.

[77] M. S. Valdés-Tresanco, M. E. Valdés-Tresanco, P. A. Valiente, and E. Moreno, “Gmx_mmpbsa: A New Tool to Perform End-State Free Energy Calculations with GROMACS,” en, Journal of Chemical Theory and Computation, vol. 17, no. 10, pp. 6281–6291, Oct. 2021. doi: 10.1021/acs.jctc.1c00645.

[78] B. R. Miller, T. D. McGee, J. M. Swails, N. Homeyer, H. Gohlke, and A. E. Roitberg, “MMPBSA.py : An Efficient Program for End-State Free Energy Calculations,” en, Journal of Chemical Theory and Computation, vol. 8, no. 9, pp. 3314–3321, Sep. 2012. doi: 10.1021/ct300418h.

[79] X. Daura, K. Gademann, B. Jaun, D. Seebach, W. F. Van Gunsteren, and A. E. Mark, “Peptide Folding: When Simulation Meets Experiment,” en, Angewandte Chemie International Edition, vol. 38, no. 1–2, pp. 236–240, Jan. 1999. doi: 10.1002/(SICI)1521-3773(19990115)38:1/2<236::AID-ANIE236>3.0.CO;2-M.

