## Supplementary information for "Conformational selection and redox-dependent destabilization in the IspG–FldA electron transfer complex"

**Table S1: Docking poses retained after conformational and inter-cofactor distance filtering.** For each docking or cofolding method, the retained IspG–FldA poses are listed with their intended IspG conformation (open, closed, or unspecified for the cofolding methods, which cannot be directed toward a particular conformational state), the inter-cofactor (FMN–[4Fe–4S]) distance, and the HADDOCK3 score (a.u.). ZDOCK generated 10 poses per conformation and HDOCK 100; for HDOCK, only those passing the 20 Å inter-cofactor distance cutoff are shown here. Lower (more negative) HADDOCK3 scores indicate more favorable poses. The top-ranked pose in each conformational state was carried forward to molecular dynamics (ZDOCK open and HDOCK closed; see main-text Figure 1).

| Method | Conformation | Cofactor dist. (Å) | HADDOCK3 score (a.u) |
|---|---|---|---|
| Boltz-2 | — | 16.2 | $-125.9$ |
| Boltz-2 | — | 15.2 | $-104.0$ |
| Boltz-2 | — | 15.7 | $-100.7$ |
| Boltz-2 | — | 15.5 | $-100.6$ |
| Boltz-2 | — | 17.1 | $-90.3$ |
| Chai-1 | — | 13.1 | $-127.8$ |
| Chai-1 | — | 14.8 | $-119.0$ |
| Chai-1 | — | 15.5 | $-118.8$ |
| Chai-1 | — | 12.9 | $-108.9$ |
| Chai-1 | — | 15.1 | $-89.9$ |
| ZDOCK | Open | 13.7 | $-104.3$ |
| ZDOCK | Open | 13.9 | $-90.3$ |
| ZDOCK | Open | 14.2 | $-77.9$ |
| ZDOCK | Open | 25.4 | $-76.5$ |
| ZDOCK | Open | 15.0 | $-75.8$ |
| ZDOCK | Open | 14.6 | $-62.9$ |
| ZDOCK | Open | 20.7 | $-61.6$ |
| ZDOCK | Open | 23.8 | $-49.3$ |
| ZDOCK | Open | 13.4 | $-49.0$ |
| ZDOCK | Open | 23.8 | $2.1$ |
| ZDOCK | Closed | 29.3 | $-95.5$ |
| ZDOCK | Closed | 29.1 | $-93.9$ |
| ZDOCK | Closed | 29.4 | $-83.6$ |
| ZDOCK | Closed | 28.7 | $-78.2$ |
| ZDOCK | Closed | 23.9 | $-73.4$ |
| ZDOCK | Closed | 23.4 | $-71.7$ |
| ZDOCK | Closed | 29.1 | $-69.7$ |
| ZDOCK | Closed | 30.8 | $-53.6$ |
| ZDOCK | Closed | 29.5 | $-44.7$ |
| ZDOCK | Closed | 28.7 | $-41.5$ |
| HDOCK | Open | 17.2 | $-80.7$ |
| HDOCK | Open | 17.5 | $-80.7$ |
| HDOCK | Open | 19.4 | $-72.2$ |
| HDOCK | Open | 18.1 | $-63.2$ |
| HDOCK | Open | 19.4 | $-52.5$ |
| HDOCK | Open | 18.3 | $-49.8$ |
| HDOCK | Closed | 18.1 | $-88.6$ |
| HDOCK | Closed | 19.5 | $-79.9$ |
| HDOCK | Closed | 13.4 | $-79.5$ |
| HDOCK | Closed | 13.1 | $-68.1$ |
| HDOCK | Closed | 20.0 | $-67.9$ |
| HDOCK | Closed | 17.9 | $-67.6$ |
| HDOCK | Closed | 14.6 | $-61.7$ |
| HDOCK | Closed | 13.2 | $-59.5$ |
| HDOCK | Closed | 17.0 | $-58.7$ |
| HDOCK | Closed | 15.0 | $-55.8$ |
| HDOCK | Closed | 14.9 | $-55.8$ |
| HDOCK | Closed | 16.4 | $-54.2$ |
| HDOCK | Closed | 13.7 | $-50.3$ |
| HDOCK | Closed | 13.9 | $-43.1$ |

**Table S2: Salt bridges at the IspG–FldA interface.** Pairs with mean occupancy $> 5\%$ across three replicates, ranked by occupancy. IspG subunit A or B is indicated in parentheses. **(a)** NHQ open: dominated by the subunit A arginine patch (R92, R106, R117, R133) engaging D135–D137 and E128/E96 of FldA, with additional subunit B contacts (R275, K325, K326). **(b)** NHQ closed: subunit B residues (R275, R351, R353, K325, K326) contact D147, D93, D67/D68, E16, D35, and E61, with R37(A) engaging D147/E92. **(c)** NSQ open: retains the subunit A arginine-patch contacts (R92, R106, R117, R133), with a shift in the dominant pairs relative to NHQ open and additional subunit B (K326) and subunit A (K181, K143) engagement.

**(a) NHQ open**

| # | Pair (IspG–FldA) | Occ. |
|---|---|---|
| 1 | R117(A)–D136 | 0.53 |
| 2 | R92(A)–D137 | 0.39 |
| 3 | R117(A)–D137 | 0.38 |
| 4 | R106(A)–D136 | 0.31 |
| 5 | R117(A)–D135 | 0.28 |
| 6 | R133(A)–E128 | 0.26 |
| 7 | R133(A)–E96 | 0.22 |
| 8 | R275(B)–D11 | 0.19 |
| 9 | R133(A)–D137 | 0.17 |
| 10 | R106(A)–E128 | 0.16 |
| 11 | R92(A)–D135 | 0.16 |
| 12 | K325(B)–E61 | 0.15 |
| 13 | R275(B)–D65 | 0.12 |
| 14 | R275(B)–D68 | 0.09 |
| 15 | R163(A)–D136 | 0.08 |
| 16 | K326(B)–D67 | 0.08 |
| 17 | K326(B)–D65 | 0.07 |
| 18 | K325(B)–D11 | 0.07 |
| 19 | R275(B)–D67 | 0.06 |
| 20 | K326(B)–D11 | 0.06 |

**(b) NHQ closed**

| # | Pair (IspG–FldA) | Occ. |
|---|---|---|
| 1 | R275(B)–D147 | 0.30 |
| 2 | R37(A)–D147 | 0.28 |
| 3 | R351(B)–D67 | 0.27 |
| 4 | R351(B)–D68 | 0.25 |
| 5 | R353(B)–D68 | 0.25 |
| 6 | R353(B)–D67 | 0.20 |
| 7 | K325(B)–D93 | 0.18 |
| 8 | R353(B)–E16 | 0.17 |
| 9 | R37(A)–E92 | 0.15 |
| 10 | R353(B)–D11 | 0.15 |
| 11 | R275(B)–D93 | 0.14 |
| 12 | R353(B)–D35 | 0.14 |
| 13 | R351(B)–E61 | 0.12 |
| 14 | K326(B)–D93 | 0.10 |
| 15 | K325(B)–E92 | 0.09 |
| 16 | R351(B)–D11 | 0.08 |
| 17 | K325(B)–E96 | 0.07 |
| 18 | R37(A)–E128 | 0.06 |
| 19 | R275(B)–D11 | 0.06 |
| 20 | R37(A)–D93 | 0.06 |

**(c) NSQ open**

| # | Pair (IspG–FldA) | Occ. |
|---|---|---|
| 1 | R133(A)–E96 | 0.54 |
| 2 | R92(A)–D137 | 0.46 |
| 3 | R106(A)–E128 | 0.45 |
| 4 | R117(A)–D137 | 0.39 |
| 5 | R117(A)–D136 | 0.27 |
| 6 | K326(B)–D11 | 0.26 |
| 7 | K181(A)–E128 | 0.24 |
| 8 | R133(A)–E128 | 0.23 |
| 9 | K326(B)–D65 | 0.22 |
| 10 | R275(B)–D11 | 0.22 |
| 11 | R37(A)–E151 | 0.16 |
| 12 | R92(A)–D135 | 0.11 |
| 13 | R275(B)–D65 | 0.10 |
| 14 | K181(A)–E96 | 0.09 |
| 15 | K325(B)–D68 | 0.08 |
| 16 | R275(B)–D67 | 0.07 |
| 17 | R275(B)–D68 | 0.07 |
| 18 | K143(A)–E61 | 0.07 |

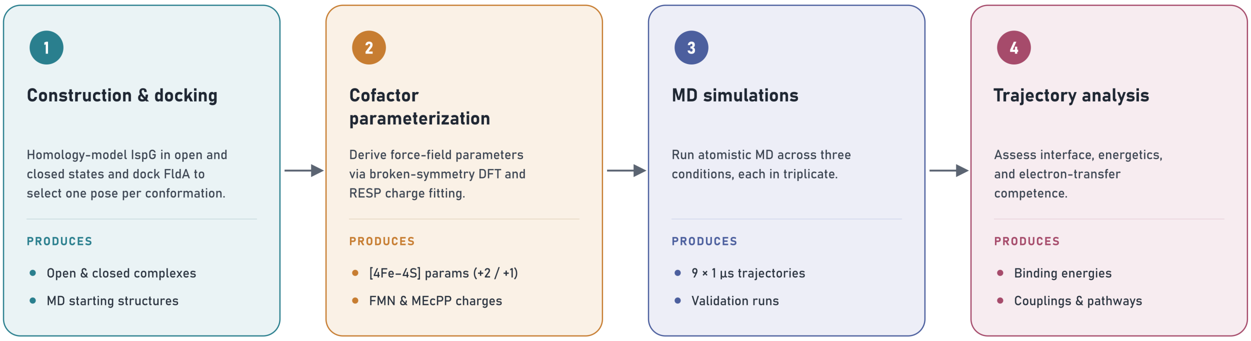


**Figure S1: Overview of the computational workflow.** The pipeline proceeds in four stages. (1) Construction and docking: *E. coli* IspG is homology-modeled in its open and closed conformations and FldA is docked onto each, yielding one selected complex per conformation as the starting structures for simulation. (2) Cofactor parameterization: force-field parameters are derived from broken-symmetry DFT and RESP charge fitting, producing custom parameters for the [4Fe–4S] cluster in both oxidation states (+2, +1) and partial charges for FMN and the MEcPP substrate. (3) Molecular dynamics: the three conditions (Table 1) are each simulated in triplicate, together with the cluster-parameter validation runs, which were singular simulations. (4) Trajectory analysis: the resulting trajectories are analyzed for interface composition and energetics, and for electron-transfer competence through electronic couplings and tunneling pathways.

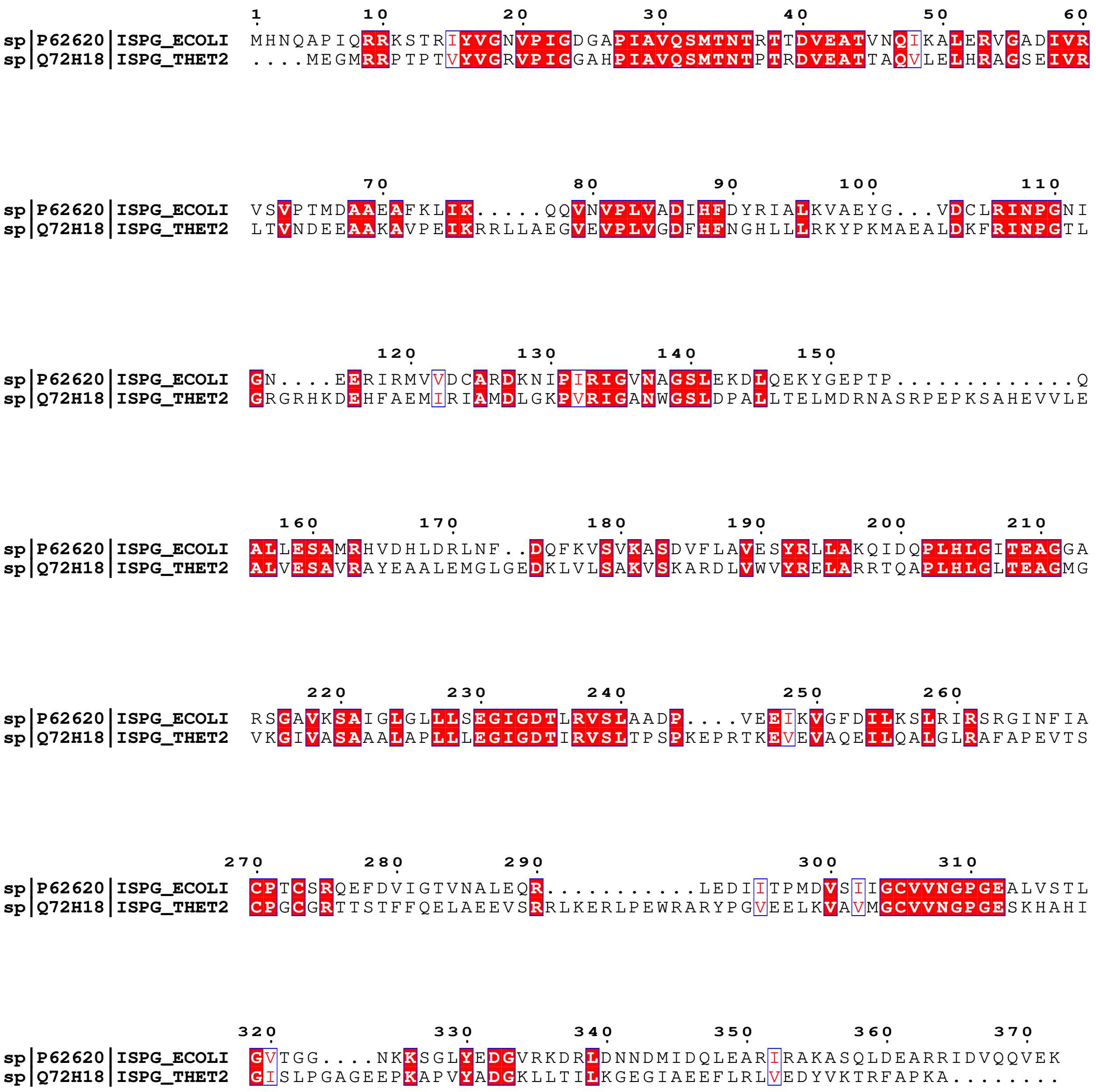


**Figure S2: Sequence alignment of *E. coli* and *T. thermophilus* IspG.** Pairwise alignment of *E. coli* IspG (UniProt P62620) and *T. thermophilus* IspG (UniProt Q72H18), the organism whose crystal structures (PDB 2Y0F, 4G9P) were used as modeling templates. The two sequences share 38% overall identity. Strictly conserved residues (identical in both sequences) are shown as white text on a red background, and similar residues as red text in white boxes; similarity was scored with the BLOSUM62 matrix. The functionally critical residues are conserved, including the three cluster-coordinating cysteines (Cys270, Cys273, Cys305 in the *E. coli* numbering), supporting the use of the *T. thermophilus* structures as references for homology modeling. Alignment generated with Clustal Omega and rendered with ESPript 3.0 (BLOSUM62 similarity scheme).

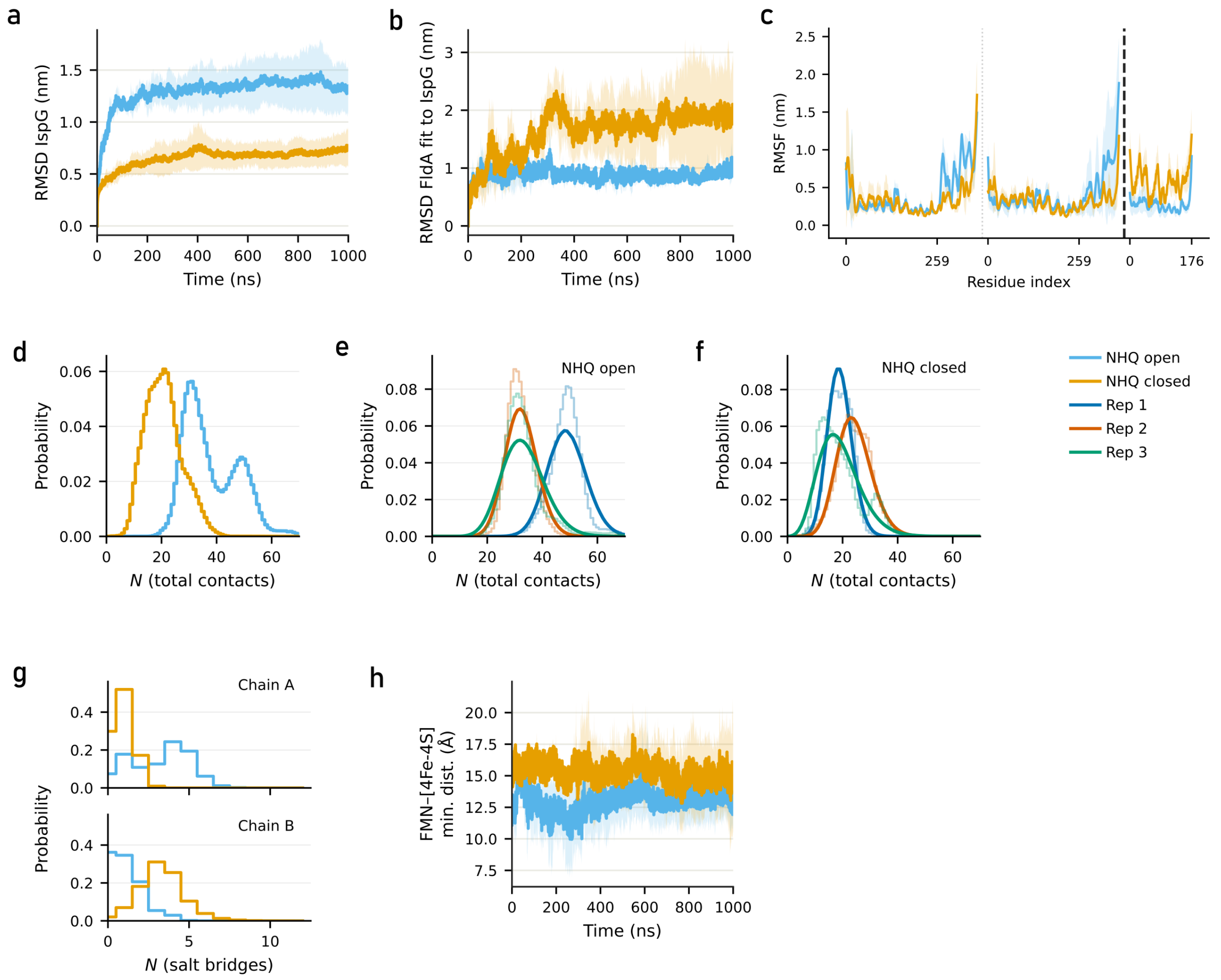


**Figure S3: Stability and interface characterization of the IspG–FldA complexes.** Data are shown for the substrate-free (NHQ open, light blue) and substrate-bound (NHQ closed, orange) complexes in the pre-electron-transfer state; where individual replicates are shown, they are colored as Rep 1 (blue), Rep 2 (red), and Rep 3 (green). (a) Backbone RMSD of IspG relative to the equilibrated starting structure as a function of simulation time, shown as the mean (solid line) and standard deviation (shaded band) across the three replicates. The IspG dimer equilibrates within the first hundred nanoseconds and remains stable thereafter in both conformations. (b) Backbone RMSD of FldA after least-squares superposition onto IspG, reporting the positional drift of FldA relative to the IspG surface. The smaller, less variable RMSD in the open complex indicates that FldA remains anchored in a narrower set of binding poses, whereas in the closed complex it explores a wider range of configurations. (c) Per-residue RMSF for the two IspG subunits and FldA (segments separated by vertical lines; residue numbering restarts per chain). (d) Distribution of the total number of IspG–FldA interface contacts (heavy-atom pairs within 4.5 Å) for the open and closed complexes, pooled across replicates. (e, f) The same total-contact distributions resolved by individual replicate for the open (e) and closed (f) complexes; light histograms show the raw per-replicate data and solid curves the corresponding density fits. (g) Distribution of the number of interface salt bridges per frame for the open (blue) and closed (orange) complexes, shown separately for IspG subunit A (top) and subunit B (bottom). (h) Minimum heavy-atom distance between the FMN cofactor and the [4Fe–4S] cluster as a function of time, shown as the mean and standard deviation across replicates. The donor–acceptor distance is consistently shorter in the open complex.

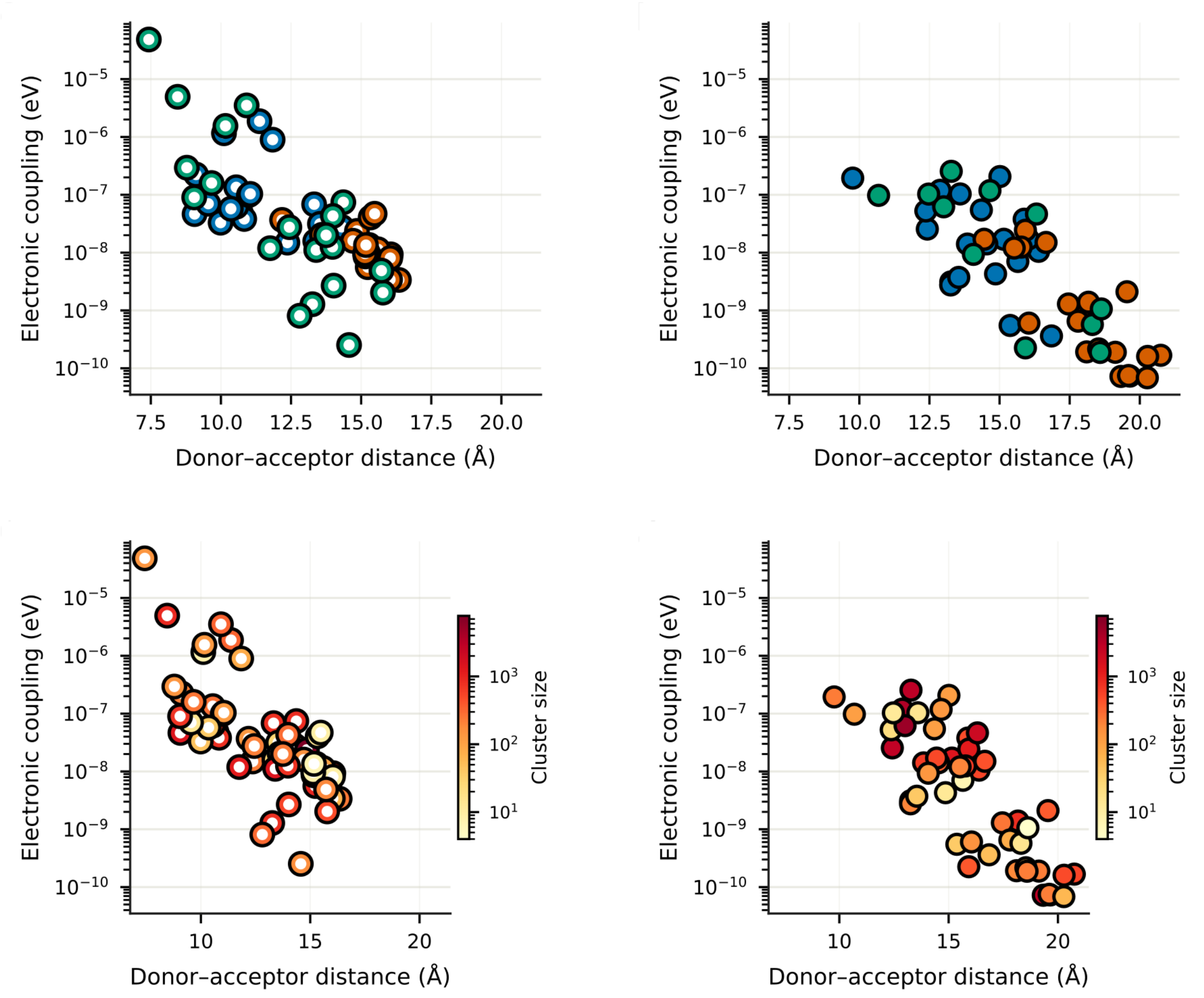


**Figure S4: Electronic coupling versus donor–acceptor distance, resolved by replicate and cluster size.** The same coupling–distance data shown in main-text Figure 3A for the open (left column: a, c) and closed (right column: b, d) IspG–FldA complexes, recolored to assess whether the coupling distribution is dominated by individual replicates or by sparsely populated conformational clusters. Each point is a representative structure from conformational clustering, with electronic coupling computed by the Pathways plugin. (a, b) Points colored by replicate (Rep 1 blue, Rep 2 red, Rep 3 green) for the open (a) and closed (b) complexes. The high-coupling configurations in the open state are sampled across multiple replicates rather than arising from a single trajectory. (c, d) The same points colored by the size (population) of the conformational cluster each structure represents, for the open (c) and closed (d) complexes. The high-coupling configurations are not restricted to small, sparsely populated clusters. Across both colorings, the open complex (a, c) reaches substantially higher peak couplings at shorter donor–acceptor distances than the closed complex (b, d).

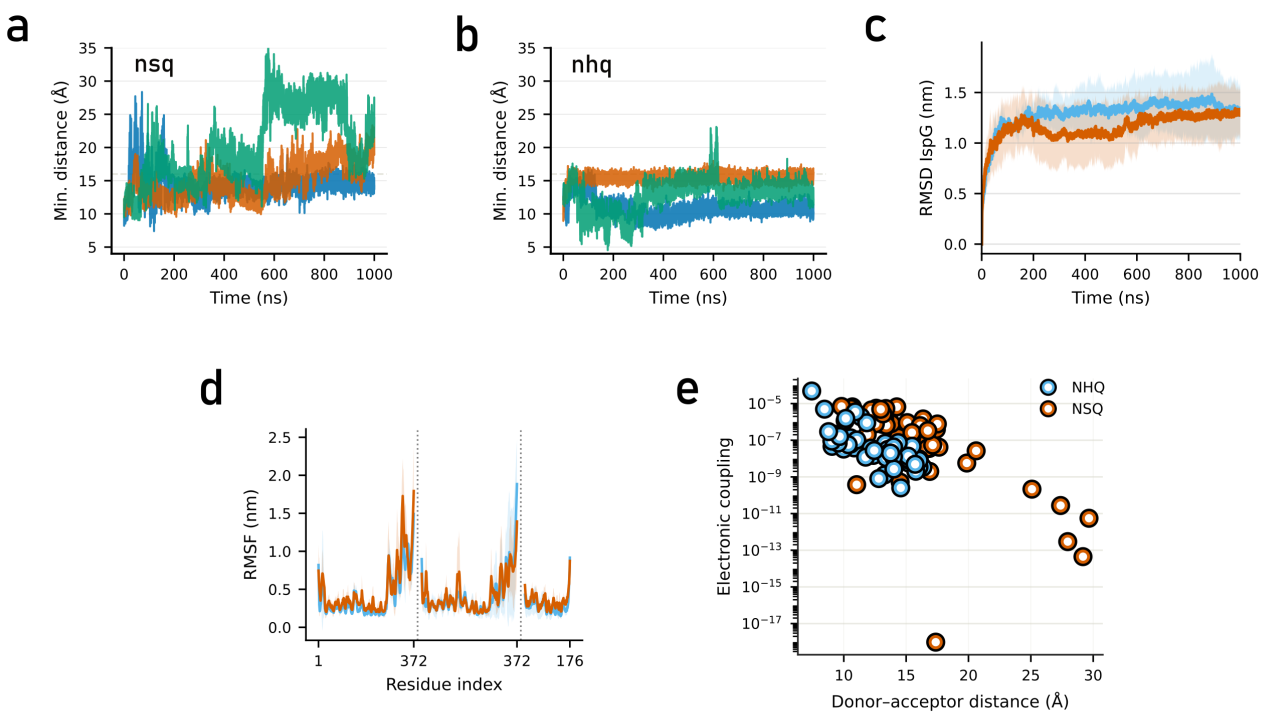


**Figure S5: Post-transfer donor–acceptor separation, structural stability, and electronic coupling of the open IspG–FldA complex.** Comparison of the pre-transfer (NHQ) and post-transfer (NSQ) states of the open *E. coli* IspG–FldA complex. (A, B) Minimum heavy-atom FMN–[4Fe–4S] distance over time for the three individual replicates (Rep 1 blue, Rep 2 orange, Rep 3 green) of the post-transfer NSQ (A) and pre-transfer NHQ (B) states. In the NSQ state the donor–acceptor distance drifts upward over the trajectory in all three replicates – modestly in two and markedly in one, which reaches ~25–30 Å after ~550 ns – whereas the NHQ replicates hold a stable separation throughout. (C) Backbone RMSD of IspG relative to the equilibrated starting structure, shown as the mean (solid line) and standard deviation (shaded band) across replicates for the NHQ (blue) and NSQ (orange) states. The fold equilibrates within the first ~200 ns and remains stable in both redox states. (D) Per-residue backbone RMSF for the two IspG subunits and FldA (segments separated by dotted lines at residue 372; numbering restarts per chain), overlaid for the NHQ (blue) and NSQ (orange) states. The flexibility profile is essentially unchanged between the two states. (E) Electronic coupling versus donor–acceptor distance for representative structures from conformational clustering, with coupling computed by the Pathways plugin. The post-transfer (NSQ, orange) state samples high-coupling ($\sim 10^{-5}$) configurations comparable to the pre-transfer (NHQ, blue) state, indicating that the transfer-competent donor–acceptor geometry does not immediately collapse on electron delivery over the timescales simulated here. This coupling is a purely geometric and electronic measure: because electron transfer shifts the relative redox potentials of the two cofactors, the retained coupling does *not* imply that reverse transfer from the reduced cluster to the oxidized FMN occurs.

#### Note S1: Conformational gating

We use the term conformational gating with two related but slightly different meanings. In its classical formulation, the term refers to a single bound complex that samples a range of orientations, only a subset of which align donor and acceptor for productive tunneling[1,2]. This is what we observe within the open IspG–FldA ensemble, where only a minority of sampled geometries achieve the peak couplings that dominate the electron transfer rate (Fig. 3A). When comparing the open and closed complexes, however, the gate operates at a coarser level: the two systems are distinct conformations of IspG rather than fluctuations within a single bound interface, and the question is whether the productive subset of geometries can be sampled at all. The two usages are complementary — the coarser gate determines which IspG conformation can host a productive interface, while the classical gate operates within that interface.

### Note S2: Kinetic comparison of substrate binding versus oxidative damage to the reduced [4Fe–4S] cluster

The claim that productive MEcPP binding outpaces oxidative damage of the post-transfer cluster rests on a pseudo-first-order comparison of substrate-binding and ROS-attack rates at physiological concentrations.

#### Oxidant rate constants for exposed [4Fe–4S] clusters

No direct measurement exists for the IspG cluster itself, so we extrapolate from the aconitase-family dehydratases, which possess a catalytically exposed Fe site structurally analogous to the non-coordinated Fe4 of IspG. Reported second-order rate constants for these systems are: superoxide, $\sim 10^6$–$10^7$ $M^{-1}$ $s^{-1}$[3,4]; hydrogen peroxide, $\sim 10^3$–$10^4$ $M^{-1}$ $s^{-1}$[5]; molecular oxygen, $\sim 10^3$ $M^{-1}$ $s^{-1}$ for well-exposed clusters such as FNR[6].

#### Cellular oxidant concentrations

Steady-state cytoplasmic levels in *E. coli* under aerobic growth are approximately 0.2 nM $O_2^{\bullet-}$, 100 nM $H_2O_2$, and 10–100 $\mu$M $O_2$[7,8,9].

#### MEcPP concentration and IspG kinetics

Intracellular MEcPP in *E. coli* falls in the $\sim$10–100 $\mu$M range under standard growth conditions, accumulating because IspG is typically rate-limiting in the MEP pathway[10]. Reported IspG $K_m$ for MEcPP lies in the 100–500 $\mu$M range across organisms and assays, placing the enzyme at or below half-saturation *in vivo*[11,12]. We assume a substrate on-rate of $10^5$–$10^7$ $M^{-1}$ $s^{-1}$, spanning the typical range for diffusion-influenced enzyme–substrate encounter[13].

#### Pseudo-first-order comparison

Multiplying each second-order rate constant by the relevant cellular concentration yields the pseudo-first-order rate at which each process proceeds on the exposed cluster (Table S3). Substrate binding outpaces the fastest-effective oxidant ($O_2$) by approximately two orders of magnitude under the most conservative on-rate assumption ($10^5$ $M^{-1}$ $s^{-1}$, 10 $\mu$M MEcPP) and three to four orders of magnitude under typical values. The relative kinetic insignificance of $O_2$ despite its high abundance arises from the spin-forbidden nature of one-electron $O_2$ reduction by a singlet cluster[8], which places $O_2$ rate constants many orders of magnitude below the diffusion limit; superoxide and hydrogen peroxide are not subject to this restriction, which is why their per-molecule rates are higher despite their lower cellular concentrations.

**Table S3:** Pseudo-first-order rates for substrate binding and oxidative attack on an exposed reduced [4Fe–4S] cluster, using cellular concentrations and rate constants from the references cited above. The pseudo-first-order rate is the product of the second-order rate constant and the cellular concentration of each species; the characteristic timescale is its reciprocal.

| Process | $k$ ($M^{-1}$ $s^{-1}$) | Concentration | Rate $k$ [conc.] ($s^{-1}$) | Timescale |
|---|---|---|---|---|
| MEcPP binding | $10^5$–$10^7$ | 50 $\mu$M | 5–500 | ms–s |
| $O_2$ attack | $10^3$ | 50 $\mu$M | $\sim 5 \times 10^{-2}$ | $\sim$20 s |
| $H_2O_2$ attack | $10^3$–$10^4$ | 100 nM | $\sim 10^{-3}$ | $\sim$15 min |
| $O_2^{\bullet-}$ attack | $10^6$–$10^7$ | 0.2 nM | $\sim 2 \times 10^{-3}$ | $\sim$10 min |

#### Caveats

(i) The cluster oxidation rate constants are extrapolated from structurally analogous but non-identical Fe–S enzymes, and no direct measurement exists for IspG. (ii) The argument requires that substrate-triggered domain closure proceeds on a timescale comparable to substrate binding itself, which has not been directly measured for IspG. (iii) Our $\mu$s simulations establish the accessibility of the outward-drifted cluster geometry, not its occupancy on biological timescales; the two- to four-order-of-magnitude kinetic margin reported here is large enough to accommodate substantial uncertainty in this occupancy.

### Supplementary Methods

#### Structure prediction and dimer construction

Monomeric *E. coli* IspG models were generated using five approaches: AlphaFold3, Boltz-2, Chai-1, Robetta, and MODELLER-based homology modeling. AlphaFold3, Chai-1, and Robetta were run via their respective web servers. Boltz-2 and Robetta were run both with and without the *T. thermophilus* IspG crystal structures 2Y0F (substrate-free) and 4G9P (substrate-bound) as templates. AlphaFold3, Chai-1, and Boltz-2 were additionally run with two copies of the *E. coli* IspG sequence to directly predict homodimeric assemblies. Chai-1 and Boltz-2 support explicit [4Fe–4S] cluster modeling, which was enabled during prediction. Because AI-based methods generate a single lowest-energy prediction without conformation-specific guidance, they cannot reliably be directed toward a particular conformational state. All predicted structures were evaluated by C$\alpha$ RMSD against the *T. thermophilus* reference structures using the CE-align algorithm in PyMOL 3 (`cealign`, default parameters). MODELLER-based homology modeling reproduced both conformations most accurately and was used for all subsequent simulations.

Homology models of *E. coli* IspG in the substrate-free and substrate-bound states were built with MODELLER 10.8[14] using the 2Y0F and 4G9P crystal structures as templates, respectively. Templates were reduced to a single chain prior to modeling. A base alignment was generated with MODELLER's `align2d` procedure and refined using SALIGN with gap opening and extension penalties of $-450$ and $-50$, and feature weights $(1.0, 0.5, 0.5, 0.5, 0.2, 0.0)$. Three independent models were generated per state using the `automodel` class with slow optimization scheduling, no MD refinement (`md_level = none`), 300 variable optimization iterations, and a deviation parameter of 9. All models were evaluated by CE-align C$\alpha$ RMSD; because values were highly similar across models, the final monomer was selected on the basis of the lowest DOPE score.

Substrate-free and substrate-bound IspG monomers were assembled into homodimers by aligning two copies of each monomer to the corresponding crystallographic dimer (2Y0F and 4G9P, respectively), generated in PyMOL by symmetry expansion (`symexp`). The [4Fe–4S] clusters and the substrate MEcPP were placed based on the same crystallographic entries, copying these objects over from the respective crystallographic entries after alignment.

The crystal structure of FldA (PDB 1AHN) was used as the structural template. Experimentally unresolved C-terminal residues were modeled using MODELLER: the target sequence was aligned to the crystal structure with `align2d`, and a single model was generated with `md_level = refine.very_fast`.

#### Docking and complex selection

Rigid-body docking of FldA onto IspG was performed with ZDOCK[15] and HDOCK[16] via their respective web servers, using the homology-modeled IspG structures as the receptor. Docking was performed separately for the substrate-free and substrate-bound IspG conformations, generating 10 poses per conformation for ZDOCK and 100 poses per conformation for HDOCK. Cofolding-based complex prediction was additionally performed with Boltz-2[17] and Chai-1[18]; these methods do not permit explicit specification of the IspG conformational state and were applied without conformational constraints.

Poses were filtered in three successive steps: (i) retention of poses matching the target IspG conformation (open or closed), assessed by C$\alpha$ RMSD against the reference structures; (ii) retention of poses with an inter-cofactor (FMN–[4Fe–4S]) distance below 20 Å; and (iii) ranking by HADDOCK3 score[19], a linear combination of van der Waals interactions, electrostatics, desolvation energy, ambiguous interaction restraints, and buried surface area. The top-ranked retained pose per conformational state, defined as the pose with the most favorable HADDOCK3 score, was used as the starting structure for molecular dynamics simulations.

#### [4Fe–4S] cluster parameterization

**QM cluster model.** The IspG [4Fe–4S] cluster is ligated by only three cysteine residues (Cys270, Cys273, Cys305), leaving one iron site without a protein ligand — a non-canonical coordination environment not covered by existing force-field parameterizations. A quantum-mechanical cluster model was constructed comprising the [4Fe–4S] core and the side chains of the three coordinating cysteines, truncated at the C$\beta$–C$\alpha$ bond and capped with hydrogen atoms. Two redox states were modeled: $[4Fe–4S]^{2+}$ (net charge $-1$, singlet multiplicity) and $[4Fe–4S]^{1+}$ (net charge $-2$, doublet multiplicity). All three coordinating cysteines were treated as thiolates (net charge $-1$ each).

**Broken-symmetry DFT calculations.** Broken-symmetry geometry optimizations were performed in Gaussian 16 at the UB3LYP[20,21] level with a mixed basis set (6-31G*[22,23] for non-metal atoms; LANL2DZ[24,25,26] for Fe), using quadratic SCF convergence, damping, an ultrafine integration grid, and a mixed initial guess. Harmonic frequency calculations at the same level confirmed the absence of imaginary frequencies. Force constants for bonded interactions involving the Fe–S cluster were derived from the frequency calculations using the Seminario method[27] as implemented in MCPB.py[28]. The stability of the broken-symmetry electronic solutions was verified with `Stable=Opt` on the converged wavefunctions, and robustness to the initial guess was confirmed by repeating single-point calculations with different initializations, all of which converged to the same solution.

**RESP charge derivation.** Partial charges for the cluster and its coordinating cysteines were derived following the RESP methodology[29]. Electrostatic potential grids were computed at the UB3LYP/6-31G*/LANL2DZ level using Merz–Kollman population analysis with MK radii (Fe radius set to 2.0 Å) and ESP grid regeneration (`Pop=(MK,ReadRadii)`, `IOp(6/33=2,6/42=6)`). RESP fitting was carried out with the `resp` program from AmberTools25[30] in a single-stage fit (`ihfree=0`, `qwt=0.0005`). No equivalencing constraints were applied to iron or sulfur atoms, preserving the asymmetric charge distribution arising from the broken-symmetry electronic state[31]. The net charge of the fitted fragment was constrained to the formal integer charge of the full cluster–cysteine unit: because each of the three coordinating cysteines contributes a $-1$ thiolate charge, the fragment charges are $-1$ for the $[4Fe–4S]^{2+}$ state (cluster charge $+2$, three thiolates $-3$) and $-2$ for the $[4Fe–4S]^{1+}$ state (cluster charge $+1$, three thiolates $-3$), with an additional electron for the reduced state. ESP grids were extracted from the Gaussian output using `espgen`; the `.ac` file was corrected prior to fitting to assign MCPB.py atom types to the iron (M1–M4) and bridging sulfide atoms (Y2–Y4, Y6), and to add Fe–$S_{cys}$ coordination bonds.

**Assembled cluster–cysteine unit.** Separate `mol2` files were generated for the [4Fe–4S] core (residue name FEO for $[4Fe–4S]^{2+}$; FER for $[4Fe–4S]^{1+}$) and for each coordinating cysteine. For the coordinating cysteines, C$\beta$ and S$\gamma$ charges were taken from the RESP fit, while all backbone atoms retained their standard ff19SB charges from the CYX residue template[32]. MCPB.py atom types were assigned to the coordinating S$\gamma$ atoms (Y5, Y1, Y7 for Cys270, Cys273, Cys305 respectively) and to the C$\alpha$ atoms (XC).

Because the QM model comprised cysteine fragments truncated at the C$\alpha$ terminus, 18 of the 32 RESP-fitted atoms are capping atoms discarded during assembly. The assembled unit therefore does not automatically sum to the formal target charge. A uniform correction was distributed equally over the 14 RESP-derived atoms retained in the final `mol2` files (four irons, four bridging sulfides, and C$\beta$/S$\gamma$ of each cysteine): $-0.039e$ per atom for $[4Fe–4S]^{2+}$ and $-0.036e$ per atom for $[4Fe–4S]^{1+}$, yielding final total charges of $-1$ and $-2$ respectively. These corrections are small relative to the magnitude of the fitted charges and are consistent with the uniform scaling approach used in analogous cofactor parameterizations[33,34].

The parameters are defined in conjunction with the coordinating cysteine residues and reproduce the correct formal charge only when used together as an assembled unit. For the $[4Fe–4S]^{2+}$ state: `FEO_core.mol2`, `C1O.lib`, `C2O.lib`, `C3O.lib`, and `FeS_twoplus.frcmod`. For the $[4Fe–4S]^{1+}$ state: `FER_core.mol2`, `C1R.lib`, `C2R.lib`, `C3R.lib`, and `FeS_oneplus.frcmod`. All parameter files are provided as Supplementary Data.

#### FMN and MEcPP parameterization

RESP charges were derived for all five FMN redox states (NHQ, AHQ, NSQ, ASQ, OX) using GAFF2 atom types. Geometry optimizations were performed in Gaussian 16 at the B3LYP/6-31G* level, followed by RESP charge fitting at the HF/6-31G* level using the `resp` program from AmberTools25[30]. For the NHQ and AHQ states, gas-phase optimizations exhibited proton migration between a ribityl hydroxyl oxygen and a nearby phosphate oxygen; these states were therefore re-optimized in an implicit diethyl ether solvent model (`SCRF=(SMD,Solvent=DiethylEther)`). Residual proton relocation persisted for the AHQ state; in this case the chemically correct protonation pattern was enforced manually by reassigning the proton (H17) from the phosphate oxygen to the ribityl oxygen (O5) and updating the corresponding bond definitions and atom types prior to RESP fitting.

MEcPP was parameterized using the same GAFF2/RESP workflow (B3LYP/6-31G* geometry optimization in Gaussian 16; HF/6-31G* RESP fitting in AmberTools25), with both phosphate groups treated as fully deprotonated (net charge $-2$). For all cofactors, RESP-ready `mol2` files were generated from the Gaussian 16 output using Antechamber with GAFF2 atom types and the appropriate net charge; missing force-field parameters were identified and generated with Parmchk2.

Of the five parameterized FMN states, NHQ and NSQ were used in production simulations. NHQ is the fully reduced, two-electron form of FldA competent for electron transfer, while NSQ is the one-electron-oxidized neutral semiquinone expected following electron delivery to IspG.

#### Electron transfer theory

Electron transfer rates in the nonadiabatic (weak-coupling) regime are described by Marcus theory[35,36]:

$$k_{\text{ET}} = \frac{2\pi}{\hbar}|H_{\text{DA}}|^2 \frac{1}{\sqrt{4\pi\lambda k_{\text{B}}T}} \exp\left[-\frac{(\Delta G + \lambda)^2}{4\lambda k_{\text{B}}T}\right] \quad (1)$$

where $H_{\text{DA}}$ is the electronic coupling between donor and acceptor states, $\lambda$ is the reorganization energy (comprising inner-sphere bond-length and angle changes at the redox centers, and outer-sphere protein and solvent polarization), and $\Delta G$ is the reaction free energy. Interprotein electron transfer at donor–acceptor separations above ~10 Å universally satisfies the nonadiabatic condition $|H_{\text{DA}}| \ll k_{\text{B}}T$[37,38], so that $k_{\text{ET}} \propto |H_{\text{DA}}|^2$.

The coupling $H_{\text{DA}}$ decays exponentially with donor–acceptor distance, but the rate of decay depends on the intervening medium. Within the Pathways formalism[39,40], the coupling through a given route is expressed as a product of per-bond attenuation factors:

$$H_{\text{DA}} \propto \prod_i \varepsilon_i \quad (2)$$

where each $\varepsilon_i$ takes the value $\varepsilon_{\text{cov}} = 0.6$ for a covalent bond, $\varepsilon_{\text{H}} = 0.36$ for a hydrogen bond, and $\varepsilon_{\text{space}} = 0.5\, e^{-1.7(r-1.4)}$ for a through-space jump of length $r$ Å. A covalent pathway is therefore orders of magnitude more conductive than one forced through space or across a poorly packed interface. Because $\Delta G$ and the inner-sphere $\lambda$ are not expected to vary significantly for the same redox pair in different conformational states, differences in $k_{\text{ET}}$ between conformations reflect primarily differences in $H_{\text{DA}}$.

#### System assembly and solvation

Titratable residue protonation states were assigned at pH 7.0 using PDB2PQR (AmberTools25[30]). Protein termini were left uncapped. Each system was solvated in an OPC[41] water box with a minimum 10 Å buffer between the protein and the box edge in all directions, and neutralized with NaCl to a physiological ionic strength; no additional ions were added.

#### Equilibration and production MD

All simulations were performed with GROMACS 2024.3[42], compiled with CUDA 12.6.0 for GPU acceleration. Proteins were described with the AMBER ff19SB force field[32] together with the OPC water model and the custom [4Fe–4S] and FMN parameters described above. No position restraints were applied at any stage.

Energy minimization used the steepest-descent algorithm (tolerance 1,000 kJ mol$^{-1}$ nm$^{-1}$; maximum 50,000 steps), followed by 100 ps of NVT equilibration with the V-rescale thermostat[43] ($\tau_T = 0.1$ ps, 310 K, applied separately to protein and solvent groups), and 100 ps of NPT equilibration with the C-rescale barostat[44] (isotropic coupling, $\tau_P = 2.0$ ps, 1.0 bar, compressibility $4.5 \times 10^{-5}$ bar$^{-1}$).

Production runs were carried out in the NPT ensemble for 1,000 ns per replicate with a 2 fs timestep, using the same thermostat and barostat settings as equilibration. Covalent bonds involving hydrogen were constrained with LINCS[45] (constraint order 4). Long-range electrostatics were treated with Particle Mesh Ewald[46] (real-space cutoff 1.0 nm; PME order 4; Fourier spacing 0.16 nm). Van der Waals interactions were truncated at 1.0 nm with long-range dispersion corrections applied to both energy and pressure. Periodic boundary conditions were applied in all three dimensions. Trajectories were written in compressed XTC format at 10 ps intervals.

The three replicates of each condition shared the same starting structure and were initialized with different random velocities. Backbone RMSD time series for all production trajectories are provided in Supplementary Figures S3A and S5C.

#### Trajectory post-processing

Raw trajectories were processed with GROMACS in two steps: molecules were first made whole across periodic boundaries (`gmx trjconv -pbc whole`), followed by removal of translational jumps (`gmx trjconv -pbc nojump`). A stride-downsampled trajectory retaining every 100th frame (one frame per 1 ns) was additionally produced for computationally intensive analyses.

#### Trajectory analysis

All RMSD calculations used the post-NPT equilibrated structure as the reference. Backbone RMSD was computed for the full complex and for the FldA subunit separately using `gmx rms` (least-squares fitting and evaluation on N, C$\alpha$, C, O atoms). The RMSD of FldA after least-squares superposition onto IspG was additionally computed as a measure of FldA positional drift relative to the complex. Per-residue RMSF was computed on backbone atoms with `gmx rmsf`. The radius of gyration of IspG and FldA was monitored with `gmx gyrate`. Hydrogen bonds between IspG and FldA were counted per frame with `gmx hbond`. The solvent-accessible surface area of the [4Fe–4S] binding pocket was computed with `gmx sasa` separately for the FldA-facing and opposing faces of the cluster. The minimum heavy-atom FMN–[4Fe–4S] distance was tracked with `gmx mindist`. The relative orientation of FMN with respect to the cluster was quantified as the angle between the plane defined by the three proximal Fe atoms and the vector from their centroid to FMN N5, computed with `gmx gangle`.

**[4Fe–4S] parameter validation.** Cluster geometry stability was assessed using dedicated 100 ns validation trajectories of the substrate-bound dimer in both redox states, run with the same equilibration and production protocol as the main simulations. The minimum S$\gamma$–Fe distance for each coordinating cysteine (Cys270, Cys273, Cys305) was computed with MDAnalysis 2.10.0[47] as a measure of coordination bond integrity, using the full trajectories. The RMSD of [4Fe–4S] atoms relative to the crystal reference structure (4G9P) was tracked, after backbone superposition.

**Interface contact analysis.** Per-frame IspG–FldA contacts were quantified with a custom MDAnalysis-based Python script[47]. Contacts were classified as: total contacts (all heavy-atom pairs within 4.5 Å), salt bridges (oppositely charged residue pairs within 4.0 Å), hydrogen bonds (donor–acceptor distance $\leq 3.5$ Å, donor–hydrogen–acceptor angle $\geq 150°$, enumerated with `HydrogenBondAnalysis`), and hydrophobic contacts (apolar heavy-atom pairs within 4.5 Å). Per-frame counts were recorded as time series; residue-pair occupancies were aggregated into contact heatmaps. Salt bridges were further characterized by per-replicate occupancy distributions, and the top-ranking pairs by mean occupancy were reported.

**Conformational clustering** Representative structures for electron transfer pathway analysis were selected by conformational clustering using the GROMOS algorithm[48] as implemented in `gmx cluster` (C$\alpha$ RMSD cutoff 0.35 nm). Trajectories were least-squares fitted on C$\alpha$ atoms and subsampled to approximately 10,000 frames prior to clustering. The twenty most populated clusters were identified, and their medoid structures — defined as the frames with the lowest mean RMSD to all other cluster members — were extracted as representatives.

**MM/GBSA binding free energies.** Binding free energies for the IspG–FldA interface were estimated with the MM/GBSA approach as implemented in gmx_MMPBSA v1.6.3[49], using the final 500 frames of each trajectory. IspG was treated as the receptor and FldA as the ligand. The Onufriev–Bashford–Case generalized Born model[50] (`igb = 5`) was applied with internal and external dielectric constants of 1.0 and 78.5 respectively, and zero salt concentration (`saltcon = 0.0`). Configurational entropy contributions were not included. Per-residue energy decomposition was performed for all residues within 10 Å of the IspG–FldA interface.

**Electron transfer pathway analysis.** Electron transfer pathways between the FMN N5 atom of FldA (donor) and the non-coordinated iron Fe4 of the IspG [4Fe–4S] cluster (acceptor) were analyzed using the Pathways plugin[51] in VMD 1.9.4a53, applied to the cluster representative structures. All tunneling pathways through the intervening protein medium were measured using the plugin's default decay parameters: $\varepsilon_{\text{cov}} = 0.6$ per covalent bond, $\varepsilon_{\text{H}} = 0.36$ per hydrogen bond, and $\varepsilon_{\text{space}} = 0.5\,e^{-1.7(r-1.4)}$ per through-space jump of length $r$ Å.

### Supplementary References


[1] V. L. Davidson, "Protein Control of True, Gated, and Coupled Electron Transfer Reactions," en, *Accounts of Chemical Research*, vol. 41, no. 6, pp. 730–738, Jun. 2008. DOI: 10.1021/ar700252c.

[2] Z.-X. Liang et al., "Dynamic Docking and Electron Transfer between Zn-myoglobin and Cytochrome $b_5$," en, *Journal of the American Chemical Society*, vol. 124, no. 24, pp. 6849–6859, Jun. 2002. DOI: 10.1021/ja0127032.

[3] D. Flint, J. Tuminello, and M. Emptage, "The inactivation of Fe-S cluster containing hydro-lyases by superoxide.," en, *Journal of Biological Chemistry*, vol. 268, no. 30, pp. 22 369–22 376, Oct. 1993. DOI: 10.1016/S0021-9258(18)41538-4.

[4] P. Gardner and I. Fridovich, "Superoxide sensitivity of the Escherichia coli aconitase.," en, *Journal of Biological Chemistry*, vol. 266, no. 29, pp. 19 328–19 333, Oct. 1991. DOI: 10.1016/S0021-9258(18)55001-8.

[5] S. Jang and J. A. Imlay, "Micromolar Intracellular Hydrogen Peroxide Disrupts Metabolism by Damaging Iron-Sulfur Enzymes," en, *Journal of Biological Chemistry*, vol. 282, no. 2, pp. 929–937, Jan. 2007. DOI: 10.1074/jbc.M607646200.

[6] J. Crack, J. Green, and A. J. Thomson, "Mechanism of Oxygen Sensing by the Bacterial Transcription Factor Fumarate-Nitrate Reduction (FNR)," en, *Journal of Biological Chemistry*, vol. 279, no. 10, pp. 9278–9286, Mar. 2004. DOI: 10.1074/jbc.M309878200.

[7] L. C. Seaver and J. A. Imlay, "Alkyl Hydroperoxide Reductase Is the Primary Scavenger of Endogenous Hydrogen Peroxide in *Escherichia coli*," en, *Journal of Bacteriology*, vol. 183, no. 24, pp. 7173–7181, Dec. 2001. DOI: 10.1128/JB.183.24.7173-7181.2001.

[8] J. A. Imlay, "Cellular Defenses against Superoxide and Hydrogen Peroxide," en, *Annual Review of Biochemistry*, vol. 77, no. 1, pp. 755–776, Jun. 2008. DOI: 10.1146/annurev.biochem.77.061606.161055.

[9] J. A. Imlay, "The molecular mechanisms and physiological consequences of oxidative stress: Lessons from a model bacterium," en, *Nature Reviews Microbiology*, vol. 11, no. 7, pp. 443–454, Jul. 2013. DOI: 10.1038/nrmicro3032.

[10] F. Rohdich et al., "The deoxyxylulose phosphate pathway of isoprenoid biosynthesis: Studies on the mechanisms of the reactions catalyzed by IspG and IspH protein," en, *Proceedings of the National Academy of Sciences*, vol. 100, no. 4, pp. 1586–1591, Feb. 2003. DOI: 10.1073/pnas.0337742100.

[11] Y.-L. Liu et al., "Structure, function and inhibition of the two- and three-domain 4Fe-4S IspG proteins," en, *Proceedings of the National Academy of Sciences*, vol. 109, no. 22, pp. 8558–8563, May 2012, Number: 22. DOI: 10.1073/pnas.1121107109.

[12] J. Misra, E. L. Mettert, and P. J. Kiley, "Functional analysis of the methylerythritol phosphate pathway terminal enzymes IspG and IspH from *Zymomonas mobilis*," en, *Microbiology Spectrum*, vol. 12, no. 7, A. Yan, Ed., e04256–23, Jul. 2024, Number: 7. DOI: 10.1128/spectrum.04256-23.

[13] A. Bar-Even et al., "The Moderately Efficient Enzyme: Evolutionary and Physicochemical Trends Shaping Enzyme Parameters," en, *Biochemistry*, vol. 50, no. 21, pp. 4402–4410, May 2011. DOI: 10.1021/bi2002289.

[14] B. Webb and A. Sali, "Comparative Protein Structure Modeling Using MODELLER," en, *Current Protocols in Bioinformatics*, vol. 54, no. 1, Jun. 2016. DOI: 10.1002/cpbi.3.

[15] B. G. Pierce, Y. Hourai, and Z. Weng, "Accelerating Protein Docking in ZDOCK Using an Advanced 3D Convolution Library," en, *PLoS ONE*, vol. 6, no. 9, O. Keskin, Ed., e24657, Sep. 2011. DOI: 10.1371/journal.pone.0024657.

[16] Y. Yan, H. Tao, J. He, and S.-Y. Huang, "The HDOCK server for integrated protein–protein docking," en, *Nature Protocols*, vol. 15, no. 5, pp. 1829–1852, May 2020. DOI: 10.1038/s41596-020-0312-x.

[17] S. Passaro et al., *Boltz-2: Towards Accurate and Efficient Binding Affinity Prediction*, en, Jun. 2025. DOI: 10.1101/2025.06.14.659707.

[18] Chai Discovery et al., *Chai-1: Decoding the molecular interactions of life*, en, Oct. 2024. DOI: 10.1101/2024.10.10.615955.

[19] M. Giulini et al., "HADDOCK3: A Modular and Versatile Platform for Integrative Modeling of Biomolecular Complexes," en, *Journal of Chemical Information and Modeling*, vol. 65, no. 13, pp. 7315–7324, Jul. 2025. DOI: 10.1021/acs.jcim.5c00969.

[20] A. D. Becke, "Density-functional thermochemistry. III. The role of exact exchange," en, *The Journal of Chemical Physics*, vol. 98, no. 7, pp. 5648–5652, Apr. 1993. DOI: 10.1063/1.464913.

[21] C. Lee, W. Yang, and R. G. Parr, "Development of the Colle-Salvetti correlation-energy formula into a functional of the electron density," en, *Physical Review B*, vol. 37, no. 2, pp. 785–789, Jan. 1988. DOI: 10.1103/PhysRevB.37.785.

[22] W. J. Hehre, R. Ditchfield, and J. A. Pople, "Self—Consistent Molecular Orbital Methods. XII. Further Extensions of Gaussian—Type Basis Sets for Use in Molecular Orbital Studies of Organic Molecules," en, *The Journal of Chemical Physics*, vol. 56, no. 5, pp. 2257–2261, Mar. 1972. DOI: 10.1063/1.1677527.

[23] P. C. Hariharan and J. A. Pople, "The influence of polarization functions on molecular orbital hydrogenation energies," en, *Theoretica Chimica Acta*, vol. 28, no. 3, pp. 213–222, 1973. DOI: 10.1007/BF00533485.

[24] P. J. Hay and W. R. Wadt, "*Ab initio* effective core potentials for molecular calculations. Potentials for the transition metal atoms Sc to Hg," en, *The Journal of Chemical Physics*, vol. 82, no. 1, pp. 270–283, Jan. 1985. DOI: 10.1063/1.448799.

[25] P. J. Hay and W. R. Wadt, "*Ab initio* effective core potentials for molecular calculations. Potentials for K to Au including the outermost core orbitals," en, *The Journal of Chemical Physics*, vol. 82, no. 1, pp. 299–310, Jan. 1985. DOI: 10.1063/1.448975.

[26] W. R. Wadt and P. J. Hay, "*Ab initio* effective core potentials for molecular calculations. Potentials for main group elements Na to Bi," en, *The Journal of Chemical Physics*, vol. 82, no. 1, pp. 284–298, Jan. 1985. DOI: 10.1063/1.448800.

[27] J. M. Seminario, "Calculation of intramolecular force fields from second-derivative tensors," en, *International Journal of Quantum Chemistry*, vol. 60, no. 7, pp. 1271–1277, 1996. DOI: 10.1002/(SICI)1097-461X(1996)60:7<1271::AID-QUA8>3.0.CO;2-W.

[28] P. Li and K. M. Merz, "MCPB.py: A Python Based Metal Center Parameter Builder," en, *Journal of Chemical Information and Modeling*, vol. 56, no. 4, pp. 599–604, Apr. 2016, Number: 4. DOI: 10.1021/acs.jcim.5b00674.

[29] C. I. Bayly, P. Cieplak, W. Cornell, and P. A. Kollman, "A well-behaved electrostatic potential based method using charge restraints for deriving atomic charges: The RESP model," en, *The Journal of Physical Chemistry*, vol. 97, no. 40, pp. 10 269–10 280, Oct. 1993. DOI: 10.1021/j100142a004.

[30] D. A. Case et al., "AmberTools," en, *Journal of Chemical Information and Modeling*, vol. 63, no. 20, pp. 6183–6191, Oct. 2023. DOI: 10.1021/acs.jcim.3c01153.

[31] L. Noodleman, "Valence bond description of antiferromagnetic coupling in transition metal dimers," en, *The Journal of Chemical Physics*, vol. 74, no. 10, pp. 5737–5743, May 1981. DOI: 10.1063/1.440939.

[32] C. Tian et al., "ff19SB: Amino-Acid-Specific Protein Backbone Parameters Trained against Quantum Mechanics Energy Surfaces in Solution," en, *Journal of Chemical Theory and Computation*, vol. 16, no. 1, pp. 528–552, Jan. 2020. DOI: 10.1021/acs.jctc.9b00591.

[33] A. T. P. Carvalho and M. Swart, "Electronic Structure Investigation and Parametrization of Biologically Relevant Iron–Sulfur Clusters," en, *Journal of Chemical Information and Modeling*, vol. 54, no. 2, pp. 613–620, Feb. 2014, Number: 2. DOI: 10.1021/ci400718m.

[34] L. Yang, Å. A. Skjevik, W.-G. Han Du, L. Noodleman, R. C. Walker, and A. W. Götz, "Water exit pathways and proton pumping mechanism in B-type cytochrome c oxidase from molecular dynamics simulations," en, *Biochimica et Biophysica Acta (BBA) - Bioenergetics*, vol. 1857, no. 9, pp. 1594–1606, Sep. 2016. DOI: 10.1016/j.bbabio.2016.06.005.

[35] R. Marcus and N. Sutin, "Electron transfers in chemistry and biology," en, *Biochimica et Biophysica Acta (BBA) - Reviews on Bioenergetics*, vol. 811, no. 3, pp. 265–322, Aug. 1985. DOI: 10.1016/0304-4173(85)90014-X.

[36] R. A. Marcus, "Electron transfer reactions in chemistry. Theory and experiment," en, *Reviews of Modern Physics*, vol. 65, no. 3, pp. 599–610, Jul. 1993. DOI: 10.1103/RevModPhys.65.599.

[37] J. J. Hopfield, "Electron Transfer Between Biological Molecules by Thermally Activated Tunneling," en, *Proceedings of the National Academy of Sciences*, vol. 71, no. 9, pp. 3640–3644, Sep. 1974. DOI: 10.1073/pnas.71.9.3640.

[38] J. R. Winkler and H. B. Gray, "Electron Flow through Metalloproteins," en, *Chemical Reviews*, vol. 114, no. 7, pp. 3369–3380, Apr. 2014, Number: 7. DOI: 10.1021/cr4004715.

[39] D. N. Beratan, J. N. Onuchic, J. R. Winkler, and H. B. Gray, "Electron-Tunneling Pathways in Proteins," en, *Science*, vol. 258, no. 5089, pp. 1740–1741, Dec. 1992. DOI: 10.1126/science.1334572.

[40] J. N. Onuchic, D. N. Beratan, J. R. Winkler, and H. B. Gray, "Pathway Analysis of Protein Electron-Transfer Reactions," en, *Annual Review of Biophysics and Biomolecular Structure*, vol. 21, no. 1, pp. 349–377, Jun. 1992. DOI: 10.1146/annurev.bb.21.060192.002025.

[41] S. Izadi, R. Anandakrishnan, and A. V. Onufriev, "Building Water Models: A Different Approach," en, *The Journal of Physical Chemistry Letters*, vol. 5, no. 21, pp. 3863–3871, Nov. 2014. DOI: 10.1021/jz501780a.

[42] M. J. Abraham et al., "GROMACS: High performance molecular simulations through multi-level parallelism from laptops to supercomputers," en, *SoftwareX*, vol. 1-2, pp. 19–25, Sep. 2015. DOI: 10.1016/j.softx.2015.06.001.

[43] G. Bussi, D. Donadio, and M. Parrinello, "Canonical sampling through velocity rescaling," en, *The Journal of Chemical Physics*, vol. 126, no. 1, p. 014 101, Jan. 2007. DOI: 10.1063/1.2408420.

[44] M. Bernetti and G. Bussi, "Pressure control using stochastic cell rescaling," en, *The Journal of Chemical Physics*, vol. 153, no. 11, p. 114 107, Sep. 2020. DOI: 10.1063/5.0020514.

[45] B. Hess, H. Bekker, H. J. C. Berendsen, and J. G. E. M. Fraaije, "LINCS: A linear constraint solver for molecular simulations," en, *Journal of Computational Chemistry*, vol. 18, no. 12, pp. 1463–1472, Sep. 1997. DOI: 10.1002/(SICI)1096-987X(199709)18:12<1463::AID-JCC4>3.0.CO;2-H.

[46] U. Essmann, L. Perera, M. L. Berkowitz, T. Darden, H. Lee, and L. G. Pedersen, "A smooth particle mesh Ewald method," en, *The Journal of Chemical Physics*, vol. 103, no. 19, pp. 8577–8593, Nov. 1995. DOI: 10.1063/1.470117.

[47] N. Michaud-Agrawal, E. J. Denning, T. B. Woolf, and O. Beckstein, "MDAnalysis: A toolkit for the analysis of molecular dynamics simulations," en, *Journal of Computational Chemistry*, vol. 32, no. 10, pp. 2319–2327, Jul. 2011. DOI: 10.1002/jcc.21787.

[48] X. Daura, K. Gademann, B. Jaun, D. Seebach, W. F. Van Gunsteren, and A. E. Mark, "Peptide Folding: When Simulation Meets Experiment," en, *Angewandte Chemie International Edition*, vol. 38, no. 1-2, pp. 236–240, Jan. 1999. DOI: 10.1002/(SICI)1521-3773(19990115)38:1/2<236::AID-ANIE236>3.0.CO;2-M.

[49] M. S. Valdés-Tresanco, M. E. Valdés-Tresanco, P. A. Valiente, and E. Moreno, "Gmx_mmpbsa: A New Tool to Perform End-State Free Energy Calculations with GROMACS," en, *Journal of Chemical Theory and Computation*, vol. 17, no. 10, pp. 6281–6291, Oct. 2021. DOI: 10.1021/acs.jctc.1c00645.

[50] D. Bashford and D. A. Case, "Generalized Born Models of Macromolecular Solvation Effects," en, *Annual Review of Physical Chemistry*, vol. 51, no. 1, pp. 129–152, Oct. 2000. DOI: 10.1146/annurev.physchem.51.1.129.

[51] I. A. Balabin, X. Hu, and D. N. Beratan, "Exploring biological electron transfer pathway dynamics with the *Pathways* Plugin for VMD," en, *Journal of Computational Chemistry*, vol. 33, no. 8, pp. 906–910, Mar. 2012. DOI: 10.1002/jcc.22927.
